# Multi-tool monitoring: integrating non-invasive methods to assess vertebrate diversity and trophic complexity in terrestrial rewilding

**DOI:** 10.1101/2025.08.01.668153

**Authors:** Clare Cowgill, James D. J. Gilbert, Ian Convery, Volker B. Deecke, Graham Sellers, Lori Lawson Handley

## Abstract

1. Rewilding, the restoration of natural processes to create self-sustaining and resilient ecosystems, is an increasingly popular conservation approach. However, outcomes are often unpredictable, and effective ecological monitoring is critical for understanding impacts. Integrating environmental DNA (eDNA) metabarcoding with other non-invasive tools may help provide more comprehensive assessments of taxonomic and functional diversity and trophic complexity.
2. We applied an integrated monitoring approach at a rewilding site in Scotland, comparing vertebrate diversity detected by metabarcoding four eDNA sample types (water, soil, tree rolling, scat) with passive acoustic monitoring (PAM) and camera trapping. Each method was evaluated for taxonomic richness, community composition, functional diversity using trait-based analyses, and trophic complexity through co-occurrence networks. The comparative effectiveness of these methods is rarely explored beyond basic taxonomic diversity metrics.
3. In total, 79 vertebrate taxa were detected: PAM captured 52 taxa, eDNA 44, and camera traps 17. Of the eDNA substrates, water and tree-rolling samples had the highest richness. Each method detected unique species, with birds and bats best covered by PAM, and small mammals by eDNA. Community composition varied between methods, with eDNA substrates capturing broader communities. PAM revealed the most pronounced community differences between wooded and open habitats.
4. Tree rolling and water eDNA captured the greatest functional diversity, while PAM showed higher redundancy, detecting species with similar ecological roles. No single method captured all functional traits. Integrating eDNA methods with PAM produced the most complete trophic network. PAM provided the most complete network but missed key terrestrial interactions that were filled by eDNA.
5. *Synthesis and applications:* Our findings highlight the necessity of an integrated multi-method ‘toolkit’ for practitioners to comprehensively capture vertebrate richness, functional diversity, and trophic complexity, particularly in rewilding contexts. We recommend combining PAM with either tree rolling or water eDNA sampling as an optimal monitoring strategy, balancing their complementary strengths in taxonomic and functional coverage.

## Introduction

Rewilding can restore ecosystems and reverse biodiversity declines by encouraging wilder, diverse landscapes. It aims to restore self-sustaining ecosystems with increased resilience and complexity by letting nature take the lead (Carver et al., 2021). Approaches range from land abandonment to species translocations and actions that kick-start ecological processes, followed by minimal intervention (Perino et al., 2019).

Despite growing popularity, empirical evidence on the ecological impacts of rewilding remains limited (Hart et al., 2023). Unlike traditional restoration, rewilding can form unpredictable ecosystems over unknown timescales (Pettorelli & Bullock, 2023), requiring systematic monitoring. Effective monitoring must capture whole-ecosystem dynamics, such as trophic complexity, disturbance, and connectivity, while remaining cost-effective (Torres et al., 2018). Developing appropriate monitoring approaches is essential for assessing rewilding and biodiversity more broadly.

Recent advances in passive acoustic monitoring (PAM), environmental DNA (eDNA) and camera trapping offer scalable solutions for monitoring ecological change, while minimising disturbance. Camera traps are widely regarded as the go-to option for monitoring terrestrial mammals, while PAM shows promise for detecting sound-producing taxa (Hoefer et al., 2023).

Environmental DNA metabarcoding is a powerful tool for landscape-scale biodiversity monitoring across broad taxonomic groups (Ruppert et al., 2019), though usage in terrestrial ecosystems lags behind those in aquatic habitats (Cowgill et al., 2025). eDNA refers to genetic material shed by organisms into environmental samples such as water, soil, or air, which can be amplified and sequenced using metabarcoding with primers conserved across broad taxonomic groups to detect multiple taxa (Taberlet et al., 2012). Terrestrial eDNA sampling presents distinct challenges compared to aquatic environments due to differences in the origin, state and transport of eDNA (Newton et al., 2025). Many substrates can detect terrestrial vertebrates with eDNA, including soil, water, air, and surface swabs from leaves and tree bark (reviewed by Cowgill et al., 2025). However, there is no “one-size-fits-all”; substrate effectiveness depends on habitat type and target species ecology (van der Heyde et al., 2020).

Despite growing adoption, the relative performance of different vertebrate detection methods remains uncertain. Comparisons between eDNA metabarcoding and camera trapping show mixed results (Cowgill et al., 2025). While eDNA from water samples can outperform camera traps for mammal detections (Sales et al., 2020), results vary across contexts (Mena et al., 2021). Soil-derived eDNA may detect small mammals missed by camera traps but can perform poorly for other vertebrates (Leempoel et al., 2020). To our knowledge, no comparisons yet exist between PAM and terrestrial eDNA metabarcoding (Cowgill et al., 2025). No single method can fully capture vertebrate diversity, requiring a multi-method approach to address biases. However the optimal combination of methods remains unknown (Newton et al., 2025).

Studies using eDNA metabarcoding and PAM have primarily focused on species richness and community composition, with functional analyses remaining underexplored (Cowgill et al., 2025; Sugai et al., 2019). Functional diversity, encompassing the ecological traits within communities, is crucial for understanding ecosystem processes (Cadotte et al., 2011).

Likewise, trophic networks based on potential interactions (metawebs; Dunne, 2005) offer a framework to understand ecosystem recovery (Boyse et al., 2025). While direct observation best captures trophic dynamics, this is impractical at large scales; non-invasive methods can support analysis of larger, more complex networks (Hartig et al., 2024). However, the effectiveness of different methods for these metrics remains poorly understood for terrestrial ecosystems.

In this study, we compare three non-invasive monitoring methods; camera trapping, PAM, and eDNA metabarcoding of four substrates (soil, water, tree rolling, and scat) for assessing vertebrate diversity in a terrestrial rewilding context. We address two questions: 1) Can a single non-invasive method reliably capture the full taxonomic, functional and trophic diversity of the vertebrate community, or is a multi-method approach necessary? 2) Is eDNA metabarcoding of one substrate sufficient to capture vertebrate diversity, or is a combination needed? Our findings aim to inform best practices for scalable, holistic monitoring of terrestrial ecosystems, ultimately supporting more robust assessments of ecosystem recovery.

## Materials and methods

### Study site

Data were collected in May and June 2023 at the Natural Capital Laboratory (NCL), a 42 ha rewilding site near Loch Ness, Scotland (Fig. 1). Established in 2019, the NCL is a demonstration site for ecological restoration and natural capital monitoring. The site comprises regenerating birch woodland, conifer plantation, felled/open ground, and peatland with previous broadleaf planting. Restoration actions include native tree planting, conifer thinning, peatland rewetting, species reintroductions and deer exclusion. The NCL also serves as a test-bed for innovative monitoring techniques, including eDNA, drone surveys, and AI-based analysis.

**Fig. 1:**
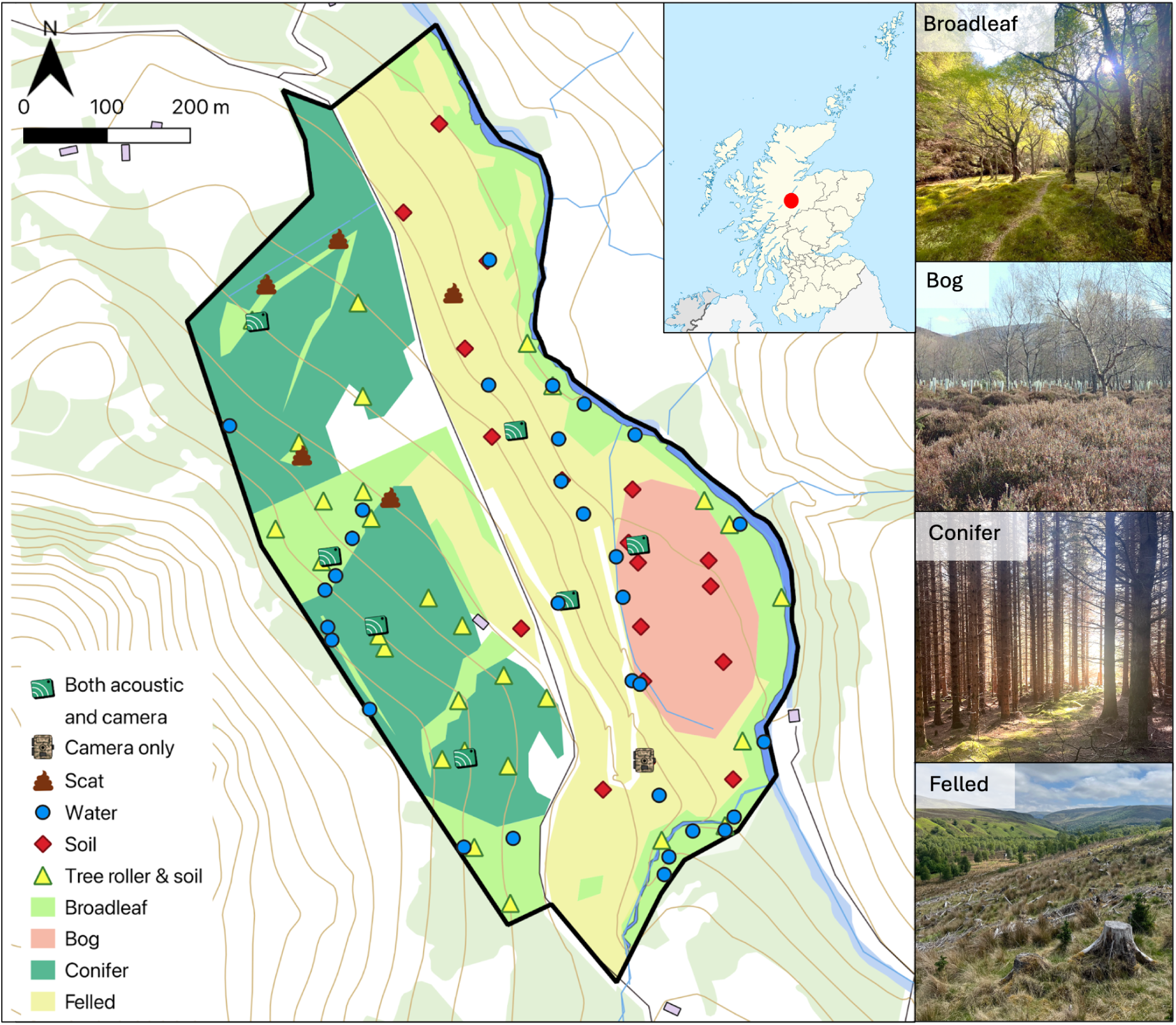
Map of the Natural Capital Laboratory rewilding site (57.1879, −4.4883) in Inverness-shire, Scotland and sampling locations. Colour identifies habitats, with eDNA sampling points and passive camera and acoustic monitoring points marked. Images show habitat examples.

### eDNA sampling

We collected 116 eDNA samples: 50 soil, 30 ‘tree roller’, 31 water, and 5 scat samples between 6 and 11 June 2023. Sampling followed stringent contamination control procedures, using sterile gloves, bleach-sterilised equipment (soaked with 10% bleach solution for 20 minutes, 5% Lipsol detergent for 5 minutes, then rinsed with purified water) and daily field blanks for water and tree roller samples. Samples were double-bagged separately according to sample type and habitat.

Water samples of approximately 1 L were collected using Whirl-Pak® bags. Due to dry weather, we adopted incidental sampling of all available waterbodies (e.g. puddles, ditches, streams). Samples were filtered on the day of collection using manifold units, through two 0.45 μm cellulose ester membrane filters (47 mm; Whatman, UK) per sample, and stored in sterile 5 ml tubes at −20 °C.

Soil sampling used stratified random sampling across habitats. At each point, 50 ml soil samples were collected from four subsamples (approximately 12.5 ml each) within a 1 m² quadrat, stored in 50 ml Falcon tubes and frozen within a few hours of collection.

Tree roller samples were taken in wooded habitats, from the same location as soil samples, following the approach of (Valentin et al., 2021) to collect DNA from the surface of tree trunks and branches. Bleach-sterilised paint rollers were wetted with purified water and rolled over tree trunks and branches for 30 seconds. The rollers were soaked in purified water, and this solution was vacuum-filtered and stored in the same way as the water samples, with one filter used per sample.

Scat samples found incidentally (n=5) were collected in 50 ml Falcon tubes and frozen within a few hours of collection.

### eDNA laboratory methods

DNA was extracted using the Mu-DNA method (Sellers et al., 2018), applying the water protocol for water/tree roller samples and soil protocol for soil/scat samples. Tree roller samples were eluted in 50 μl buffer. Soil and scat samples (2.5 g) were homogenised using a TissueLyser II (Qiagen Ltd. UK) in 10 ml jars with 5.5 ml lysis buffer. Two replicates per soil/scat sample were processed separately. Extraction blanks (buffers only) were included, with additional lysis blanks for soil/scat samples. DNA yield was assessed via Nanodrop (Thermo Fisher Scientific Ltd. UK) and extracts stored at −20 °C.

We used nested two-step metabarcoding with dual MID-tagging (Griffiths et al., 2023), targeting the vertebrate 12S mitochondrial region with primers 12S-V5-F and 12S-V5-R (Kelly et al., 2014; Riaz et al., 2011). First PCRs used Q5 High-Fidelity polymerase (Qiagen Ltd. UK) for water/tree samples and MyTaq HS (Meridian Bioscience, UK) for soil/scat.

Sub-libraries included blanks, PCR negatives, and positive controls (*Astatotilapia calliptera*, 0.05 ng/μl). Triplicate PCRs were pooled by band intensity on 2% agarose gels, and samples that showed signs of potential inhibition by poor or no amplification were diluted 1:10 and re-amplified. Amplicons were purified using magnetic bead cleanup (0.9× and 0.15× Mag-BIND^®^ beads, OmegaBiotek). Second round PCR added Illumina Unique Dual Indexes (Illumina Cambridge Ltd. UK) and underwent bead cleanup (0.7× and 0.15× Mag-BIND® beads). Sub-libraries were quantified using Qubit with a dsDNA HS Assay Kit (Thermo Fisher Scientific Ltd. UK) and qPCR (NEBNext Library Quant Kit, New England Biolabs, UK), pooled in equal concentrations and diluted to 4 nM. The final library was sequenced on an Illumina MiSeq (with 2 × 300 bp v3 chemistry) with 10% PhiX.

Raw reads were processed with a custom reproducible workflow, Tapirs (https://github.com/EvoHull/Tapirs). Sequences were trimmed, quality-filtered (Q30), and merged using fastp (Chen, 2023), then clustered with VSEARCH (Rognes et al., 2016). Taxonomic assignment used a UK vertebrate 12S database (Harper et al., 2018; see data repository), applying a majority lowest common ancestor approach (≥98% identity, ≥90% coverage; ≥80% agreement among top 2% BLAST hits).

Sample-type-specific, species-level limits of detection (LOD) were calculated based on field and lab control reads:

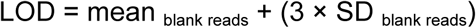

Unique LOD thresholds were applied per species and sample type. Read counts below the LOD were set to zero. Low frequency noise was removed using a 0.1% read threshold per sample. Human, domestic species (*Homo sapiens, Sus scrofa, Ovis aries, Bos taurus, Canis lupus*) and fish detections were excluded.

### PAM and camera monitoring

Two trail cameras (Browning BTC-8E-H4) facing opposite directions were deployed at eight locations. The two cameras at each location were treated independently. For comparison with eDNA sampling, data from 1st May −13th June 2023 was analysed, covering five weeks prior to and including the week of eDNA sampling. Photos were manually audited, with ambiguous records verified on iNaturalist (https://www.inaturalist.org/). Records were considered independent if the species was not spotted for 10 minutes prior, yielding 477 species records.

AudioMoth PAMs (v1.2.0; Open Acoustic Devices, UK) at the same eight locations recorded 30 seconds hourly at 192kHz. Recordings from the same time period as cameras were analysed. One device was excluded due to technical errors. Birds were identified using BirdNET Analyzer (Kahl et al., 2021) with location filtering for 193 potential species. From 6290 recordings, 10,921 verified bird identifications were obtained after manual verification of at least 20 recordings per species, per week. Small mammal calls were processed using the BTO Acoustic Pipeline (version 5.502; https://www.bto.org/our-work/science/research-areas/acoustic-monitoring), with 301 records after manual verification.

### Data analysis

A presence-absence matrix was generated for species detections in each eDNA sample or week of camera trap and PAM data for each device. Weekly aggregation helped to balance these data sets with the temporal resolution of eDNA samples, particularly given the short duration of daily recordings (12 minutes per day) and the relatively stable, non-migratory communities expected during the study period. Taxa identified at multiple taxonomic levels were consolidated at the lowest consistently detected level (see Supporting Information), with detections merged at the most precise level when both species and higher ranks were found. PAM and camera methods typically resolved taxa to species level, with higher-level identifications more common in eDNA data.

Data analysis was performed in RStudio version 4.4.2 (R Core Team, 2025); see Supplementary Table 11 for package references. Data were manipulated with ‘tidyverse’, aggregated into weekly units using ‘lubridate’, and visualised with ‘ggplot2’. Taxonomic richness was modelled using Poisson generalised linear models (GLMs) fitted with the ‘MASS’ package. Model comparisons showed method type was a stronger predictor of richness than habitat or location (Supplementary Table 8). Species accumulation curves were generated with ‘iNEXT’, for eDNA samples and daily pooled PAM and camera trap data (16 cameras over 44 days, 660 deployment days; 7 PAMs over 44 days, 308 deployment days). Community composition was analysed using non-metric multidimensional scaling (NMDS) with Jaccard dissimilarity in ‘vegan’, performed on the full dataset grouped by sampling method, and within each method to assess habitat-specific distinctions. Community differences were assessed with pairwise PERMANOVAs (‘pairwiseAdonis’).

Functional diversity was based on body mass, lifestyle, and trophic traits from published sources, with additional sources used to fill data gaps (see Supplementary Table 12).

Samples with ≥ 3 species were included in this analysis. Functional diversity was quantified using Rao’s quadratic entropy (RaoQ; the average functional difference between two species based on abundance and pairwise functional dissimilarities, Botta-Dukát, 2005) and Functional Redundancy (FRed, the difference between species diversity and RaoQ; de Bello et al., 2016). Metrics were calculated using ‘SYNCSA’, following a similar approach to Aglieri et al. (2021) for presence-absence data, using Gower’s dissimilarity of traits. Dunn tests (‘dunn.test’) compared survey methods, and trait variation was visualised using Principal Coordinates Analysis (PCoA).

Trophic complexity was assessed using a metaweb approach, constructing potential trophic interaction networks among co-occurring species by combining presence-absence data with known trophic links (Dunne, 2005). Trophic interactions from the GLOBI database were used (Poelen et al., 2014) and supplemented with additional sources (see data repositories).

Species trophic levels were calculated using the PreyAveragedTL function (‘cheddar’ package), increased by one due to the absence of primary producers. Co-occurrence of taxa was defined as detection at the same site within ±7 days for camera and PAM data, or within the same eDNA sample. Metawebs were constructed for each method, and by combining all methods, and visualised using ‘igraph’. We calculated the following metrics for the constructed networks: linkage density (mean links per taxon), connectance (proportion of realised to possible links), robustness (proportion of species removed before 50% of the network remains), modularity (strength of division into subgroups), and mean trophic level (average PreyAveragedTL). Scat samples were excluded due to small sample size and their focus on diet rather than co-occurrence, which could inflate connectance.

## Results

After quality control, eDNA data consisted of 17,517,345 paired-end reads, with an average read depth of 151,012 reads per sample. Of these, 12,074,565 reads were successfully assigned to taxa. Low-level contamination was observed in some controls and is detailed in Supplementary Table 1.

### Alpha diversity

Seventy-nine vertebrate taxa were detected across all methods. PAM detected 53 species over 308 deployment days, predicted to approach a plateau of around 70 species after 1000 days (Fig. 2A). Camera traps detected 12 species over 660 deployment days. Combining acoustic and camera data (“Both”, Fig. 2A) increased total detections to 60.

**Fig. 2:**
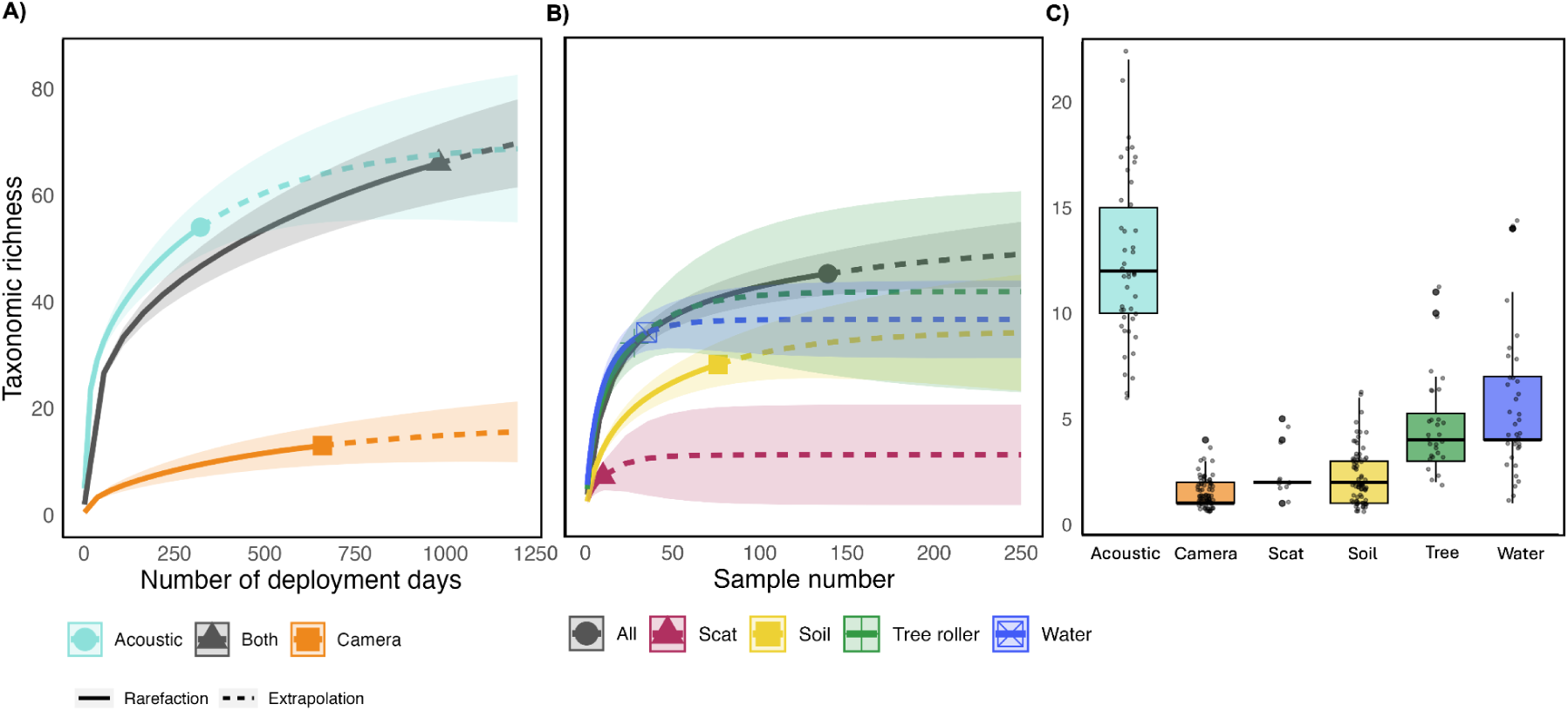
Species accumulation curves (with 95% confidence intervals) of taxonomic richness, calculated with Chao II estimator and 999 bootstrap permutations. **A)** camera trap (blue) and acoustic recorders (orange), plotted against deployment days, pooled data in grey (Both). **B)** eDNA substrates against number of samples. ’All’ refers to pooled results using all environmental substrates (soil, tree rolling and water). **C)** Taxonomic richness for each method with points equating to one sample (eDNA) or week of monitoring (other methods).

After quality control, 44 vertebrate taxa were detected across eDNA methods (Fig. 2B). Water samples showed the highest richness (34 taxa), though extrapolation predicted tree rolling could exceed this (Fig. 2B). Approximately 25 water or tree rolling samples recovered comparable richness (∼32 species) to 50 PAM days. Soil sampling detected 28 taxa, requiring 250 samples to match water sampling richness. Scat samples had the lowest richness, reflecting the low sample numbers (three *Martes martes* scats, one *Buteo buteo*, one with no detectable predator DNA). Maximum richness was achieved by combining all eDNA samples (“All”, Fig. 2B), though ∼50 water or tree roller samples showed potential to detect most vertebrates.

Median taxonomic richness differed among methods (Kruskal-Wallis: X² = 197.9, p < 0.001; Fig. 2C). PAM showed the highest median richness, while cameras showed the lowest (Fig. 2C). No significant difference was found between the median richness of water and tree rolling samples (Dunn’s test, p = 0.435), though both exceeded soil and scat sample richness (Dunn’s tests, all p < 0.016).

### Community composition

Across methods, 22 mammal, 54 bird and 3 amphibian taxa were detected (Fig. 3A). Amphibians were found only with eDNA. Of the mammals, 18 were detected with eDNA, six with cameras, and four with PAM. No mammals were detected by all three methods. Small terrestrial mammals were exclusively detected with eDNA, except common shrew (*Sorex araneus),* also detected acoustically. Five mammal taxa were detected with both eDNA and cameras, *Vulpes vulpes* only with cameras, and bats only with PAM. Of the 54 bird taxa, 48 were found with PAM (Fig. 3B). Four bird taxa were detected with all three methods, and 13 additionally with both PAM and eDNA. Four birds were exclusive to eDNA, including semi-aquatic *Cinclus cinclus*. *Accipiter gentilis* was detected with soil eDNA and cameras but not PAM.

**Fig. 3:**
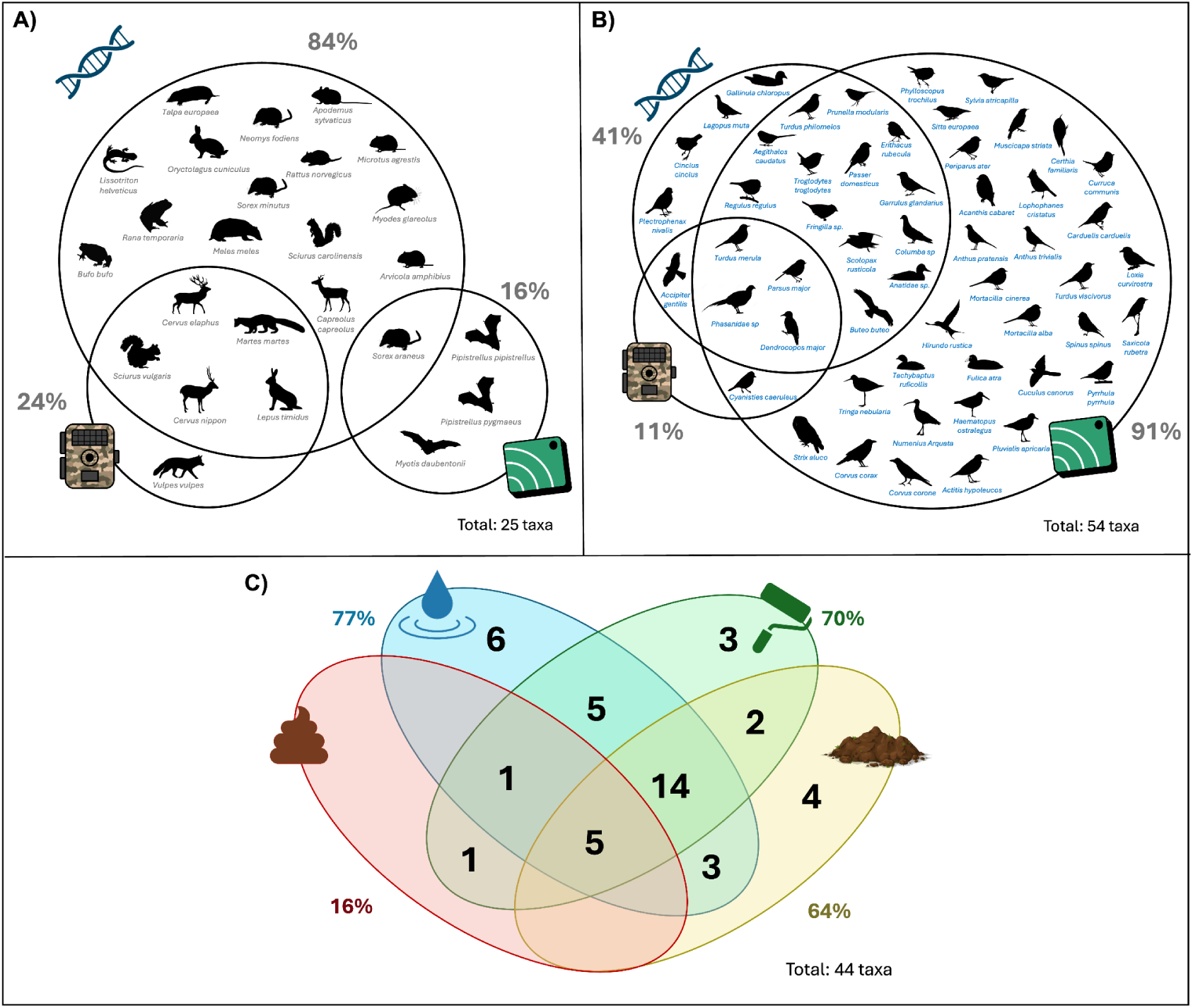
Taxa detected with eDNA sampling, camera trapping and passive acoustic monitoring split by **A)** mammals and amphibians, and **B)** birds. **C)** Number of taxa detected with each eDNA substrate and overlap. Labels show the percentage of species detected per method. Silhouettes from phylopic https://www.phylopic.org.

Of the 44 taxa detected with eDNA, five were found across all substrates, with 14 also shared across water, soil, and tree rolling (Fig. 3C). 77% of eDNA detections were found with water, 70% with tree rolling, 64% with soil and 16% with scat samples (Fig. 3C). The three environmental substrates each detected at least three unique taxa, reflecting substrate-specific habitats, e.g. *Dendrocopos major* only found with tree rolling, *Talpa europaea* with soil, and *Neomys fodiens* in water.

Communities differed between methods (PERMANOVA: R² = 0.313, p < 0.001; Fig. 4). Pairwise differences were largest between PAM and camera traps (Pairwise PERMANOVA: R² = 0.49, p=0.015), and smallest among eDNA substrates (all R² < 0.09; Supplementary Table 5). Soil samples showed the lowest pairwise differences with other methods.

**Fig. 4:**
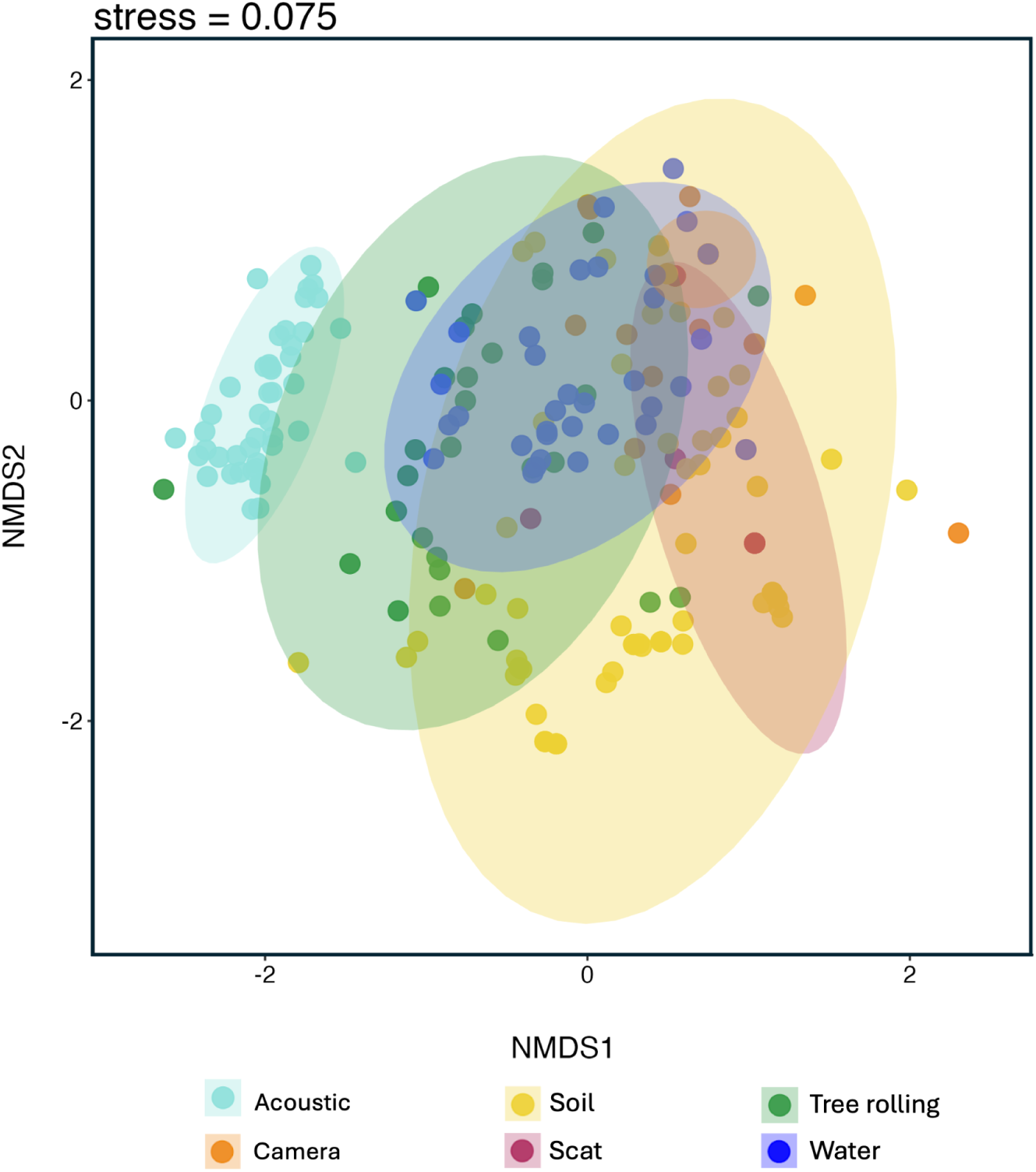
NMDS plot based on Jaccard dissimilarity of vertebrate community composition with different monitoring methods. PERMANOVA: R² = 0.318, p < 0.001. Ellipses show 95% confidence intervals.

PAM showed the strongest community distinctions between habitats (R² = 0.318, p < 0.001), revealing differences between wooded and open habitats (Fig. 5A; Supplementary Table 6).

**Fig. 5:**
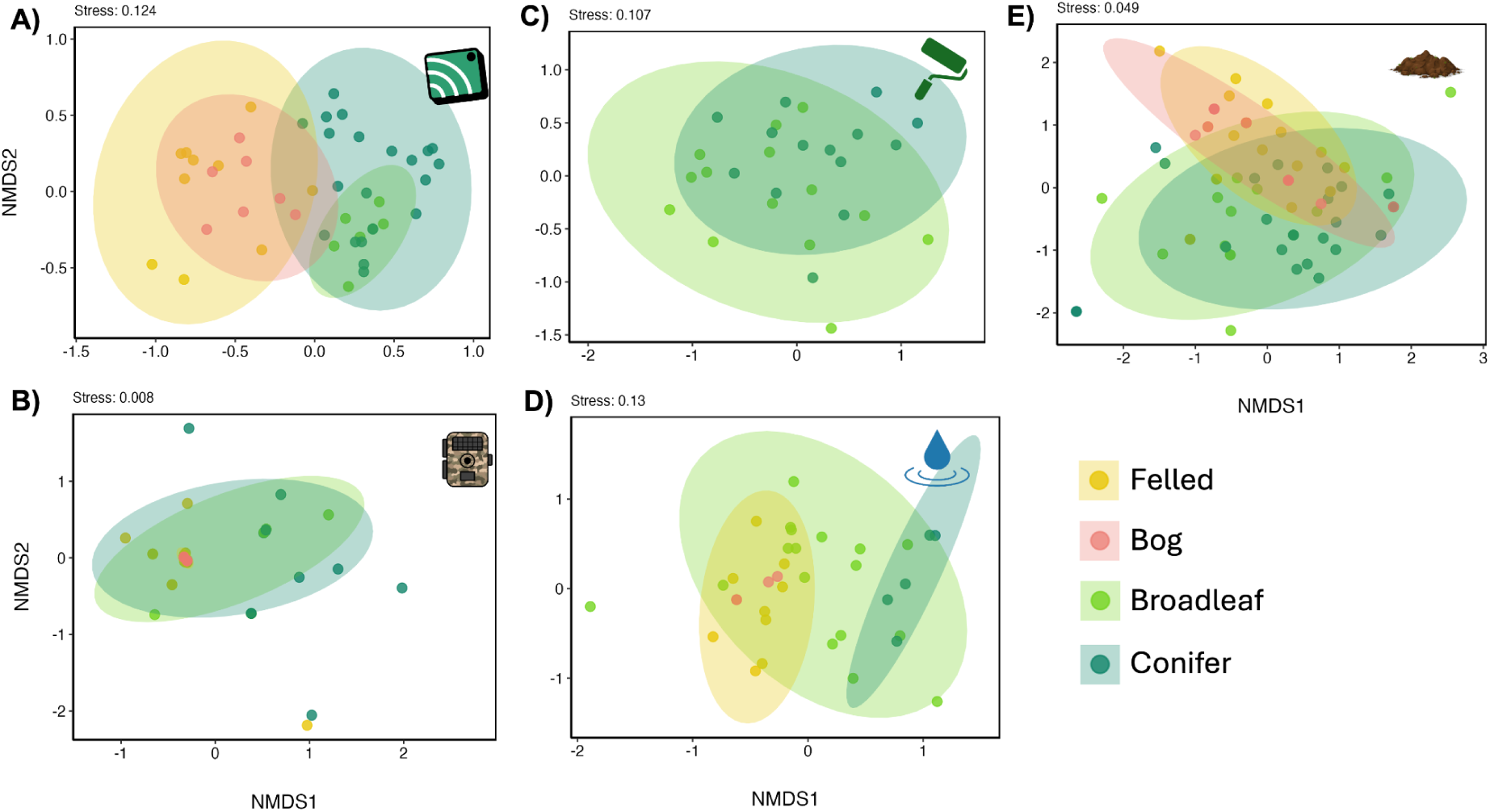
NMDS plots showing vertebrate community composition differences in habitats using Jaccard dissimilarity for each method. For passive sampling **(A-B)**, points represent weekly data per device. For eDNA (**C-E**), points represent samples.

Water samples showed distinction between conifer and felled habitats (R²=0.24, p=0.006; Fig. 5D). Soil samples revealed community differences between all wooded and non-wooded habitats (R² = 0.13, p < 0.001; Fig. 5E), reflecting the site’s spatial pattern of woodland in the west and open habitats to the east. Tree rolling was the only method that found a significant, albeit small, difference between wooded habitats (R²=0.07, p=0.007).

### Functional diversity

Functional diversity (RaoQ) differed between methods (Kruskal-Wallis: X² = 54.51, p < 0.001, Fig. 6A). Tree rolling and water samples revealed greater functional diversity than soil, PAM, or cameras (Fig. 6A; Dunn test: p < 0.008). PAM showed higher functional redundancy than other methods (Fig. 6B; Dunn test: p < 0.001), detecting many species sharing similar functions.

**Fig. 6:**
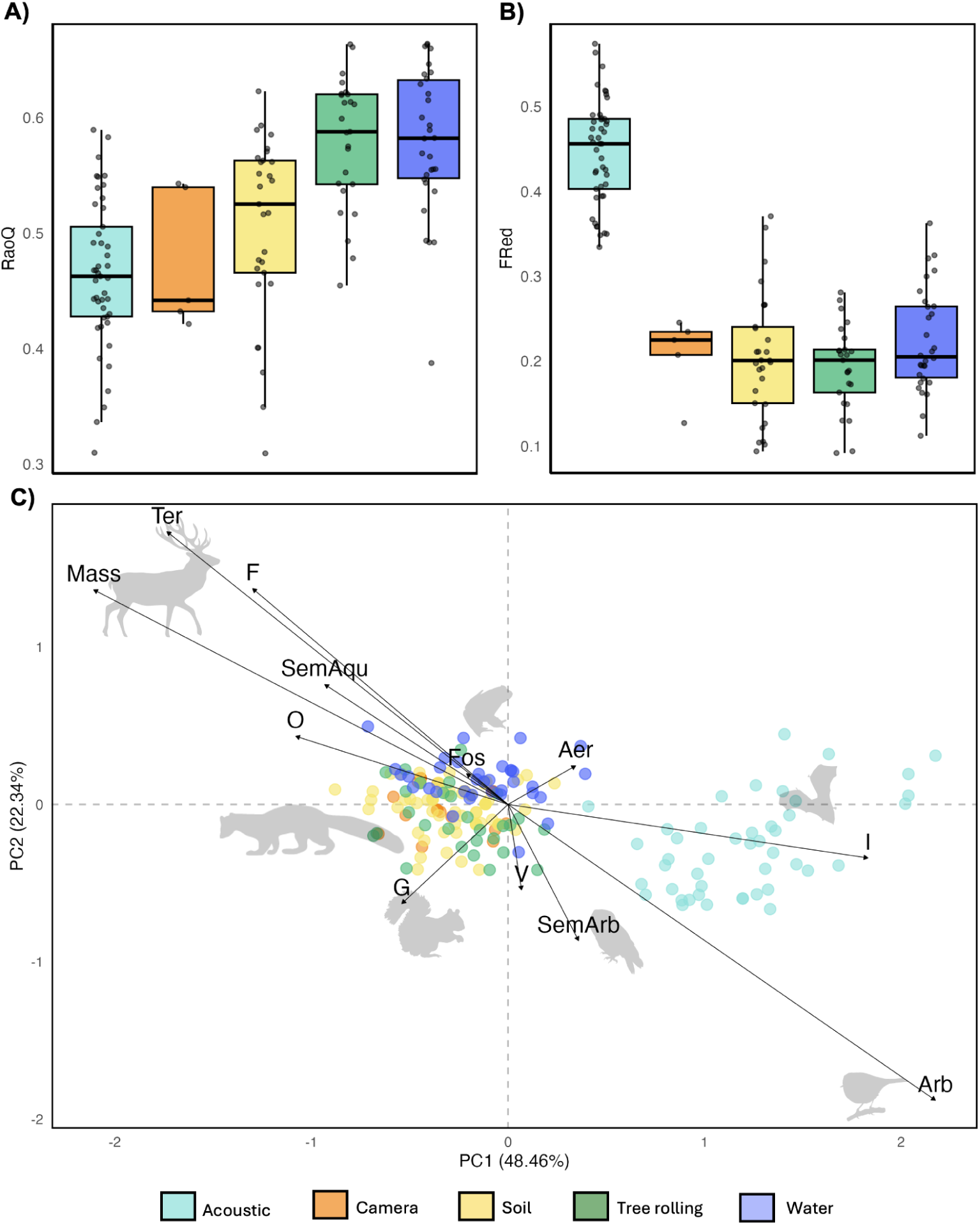
Functional diversity estimates per method: **A)** Rao’s quadratic entropy (per eDNA sample or week of passive sampling) and **B)** Functional Redundancy. **C)** PCoA of functional traits detected using different monitoring methods. Arrows show trait directions and magnitude. Functional traits: Mass (log); Trophic niche (F = Folivore, G = Granivore, O = Omnivore, I = Invertivore, V = Vertivore); Habitat type (SemAqu = Semi-Aquatic, Fos = Fossorial, Ter = Terrestrial, SemArb = Semi-Arboreal, Arb = Arboreal, Aer = Aerial). Silhouettes show species examples.

PCoA revealed that PAM detected smaller, arboreal invertivores, distinguishing this method from the others, which instead detected more larger, terrestrial folivores and omnivores (Fig. 6C). Within eDNA samples, tree rolling was associated with arboreal, granivorous taxa, while water samples were more linked with semi-aquatic taxa.

### Trophic complexity

The combined co-occurrence metaweb revealed 179 distinct trophic links (Fig. 7). PAM recovered 76% of links, weighted towards birds (Fig. 7A, 7E), while camera traps detected 3% (Fig. 7B, 7E). Combined eDNA methods recovered 29% of links, including interactions involving mammals missed by PAM (Fig. 7C). Water samples found 16% of links, tree rolling 15% and soil 6% (Fig. 7B, 7C, 7E). PAM and tree rolling together captured 89% of links, the highest coverage of any two-method combination. Adding soil or water to this pair increased coverage to over 90% and combining PAM with all eDNA substrates captured 98% of links.

**Fig. 7:**
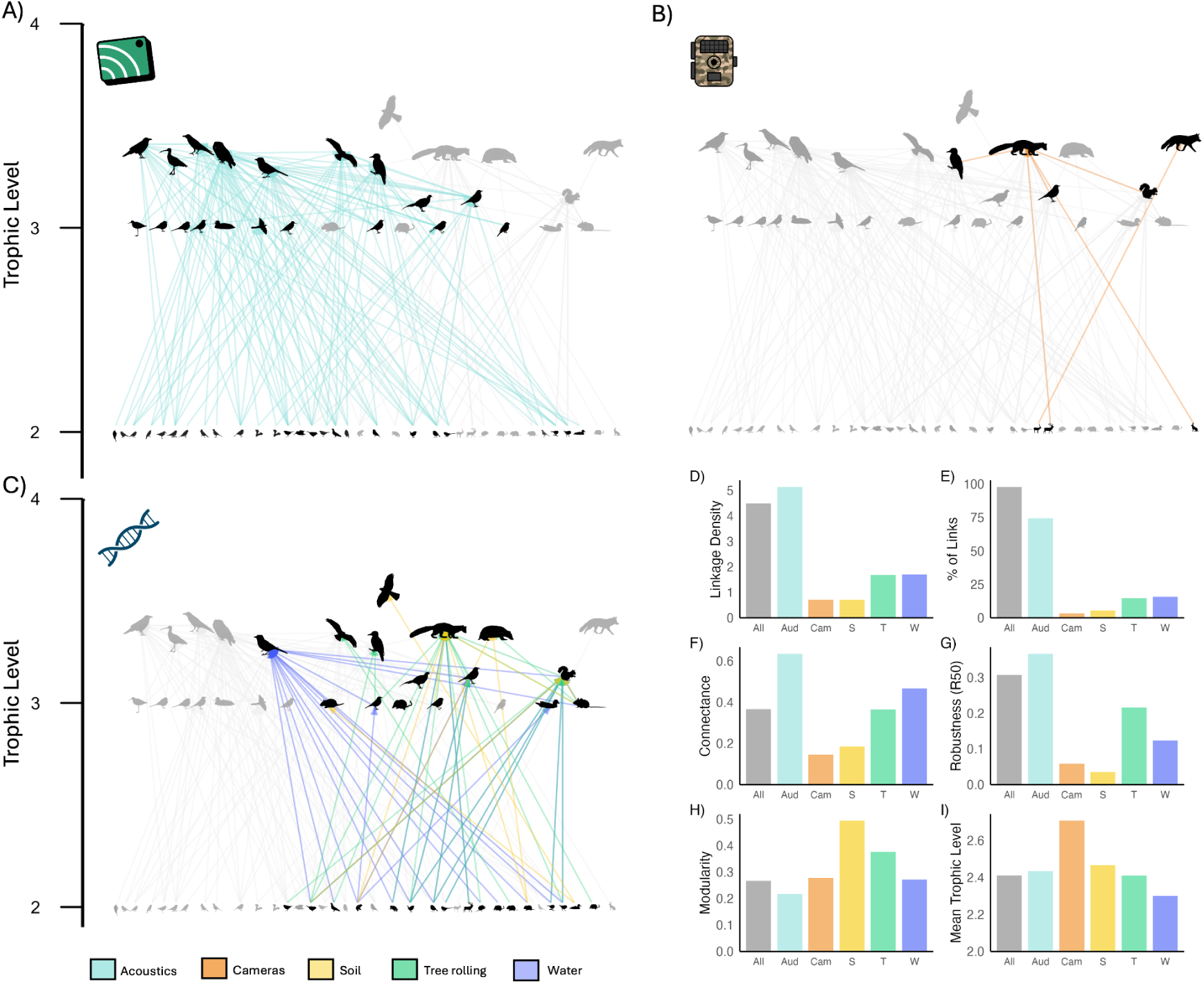
Co-occurrence metawebs of species with known trophic interactions, detected with **A)** passive acoustic monitoring, **B)** camera traps, and **C)** eDNA substrates, coloured by method type, with links found by multiple eDNA substrates shown by overlaying colours. Links and nodes not detected with the highlighted methods are grey. **D–I** show network metrics calculated for each method and all methods combined: **D)** Linkage density; **E)** Percentage of detected links; **F)** Connectance; **G)** Robustness; **H)** Modularity; **I)** Mean trophic level.

PAM showed highest linkage density and connectance, exceeding even the combined network (Fig. 7D, 7F). Acoustics and tree rolling created the most robust networks (Fig. 7G), while soil showed highest modularity, detecting relatively disconnected interactions (Fig. 7H). Cameras had a large proportion of species filling high trophic positions (Fig. 7I).

## Discussion

Non-invasive monitoring tools are increasingly used for terrestrial vertebrate surveys due to their scalability and minimal disturbance (Hartig et al., 2024). Our results show that different methods reveal complementary subsets of vertebrate biodiversity, with no single method capturing the full taxonomic and functional diversity and trophic complexity. Combining PAM with eDNA metabarcoding provided the most comprehensive assessments. Among eDNA substrates, the greatest taxonomic and functional diversity was detected with tree rolling and water samples. These findings support an integrated approach for monitoring vertebrate diversity in the context of rewilding and terrestrial monitoring in general.

### Taxonomic richness and community composition

Methods differed in terms of species detections. PAM recorded the highest taxonomic richness, with most (49/53) being birds, the most species-rich UK vertebrates (McInerny et al., 2022). PAM achieved this with comparatively low sampling effort, emphasising its efficiency for long-term monitoring. eDNA metabarcoding captured 44 taxa, including small mammals and amphibians missed by other approaches. Camera traps detected only 15% of taxa due to selectivity for large-bodied mammals, of which the UK has few species (Mathews et al., 2018), unlike in high-diversity regions where camera traps can match eDNA (Mena et al., 2021). Scat sampling, though effective for low-density species (van der Heyde et al., 2020), was limited here by sample availability. These findings align with a recent meta-analysis showing that the comparative effectiveness of eDNA metabarcoding varies across contexts for vertebrate surveys (Cowgill et al., 2025), with detections influenced by species’ body size and visitation frequency (Ryan et al., 2022). To our knowledge, this study is the first to directly compare eDNA and PAM. These methods are complementary, with PAM offering rapid detection of sound-producing species and eDNA providing coverage of less vocal or elusive taxa.

Strategic substrate selection may be more cost-effective than comprehensive multi-substrate sampling for vertebrate eDNA monitoring. Combining all eDNA substrates did not substantially increase richness beyond that potentially achieved by 50 water or tree-rolling samples, challenging the assumption that more substrates yield higher richness (Cowgill et al., 2025). Rarefaction analyses revealed non-plateauing curves across methods, indicating that no method realised its taxonomic richness potential. This supports focussing on fewer, higher-yield substrates, but note that using multiple primer regions could have enhanced taxonomic coverage (Ruppert et al., 2019).

Discriminating between habitats is important for example to track changes in community composition and connectivity. Methods varied in distinguishing vertebrate communities by habitat. PAM clearly distinguished open from wooded habitats, likely due to differences in sound propagation and community composition (Winiarska et al., 2024). Soil eDNA was also effective for distinguishing open and wooded habitats, though distinctions were smaller. Only tree rolling differentiated between coniferous and broadleaf woodland communities.

Detection capabilities could be influenced by habitat-specific factors affecting sound propagation or DNA preservation. Tree rolling is obviously limited to wooded areas, capturing DNA from species that contact tree surfaces (Allen et al., 2023), with signals likely influenced by woodland composition and structure. Water eDNA can lack fine-scale spatial specificity as DNA can originate from distant sources (Carraro et al., 2020). UV exposure degradation may be higher in open habitats, driving community distinctions (Guthrie et al., 2024). These findings show that pooling data from different methods can mask ecological patterns, as each approach may respond differently to habitat variation and thus contribute unique information that could be lost if combined without consideration.

### Functional Diversity

High taxonomic richness does not necessarily reflect functional richness, particularly in systems with high redundancy in ecological roles. Ecosystem functioning is driven by traits of abundant or high-biomass taxa (Smith et al., 2020), and methods missing functionally dominant species may fail to represent functional complexity. Tree rolling and water eDNA sampling captured the highest functional diversity, despite PAM detecting more species.

These methods sample across niches: tree rolling captures arboreal to terrestrial taxa (Allen et al., 2023); water integrates DNA across spatial scales via runoff (Carraro et al., 2020), detecting terrestrial and aquatic taxa. This echoes findings from aquatic systems, where eDNA outperforms traditional methods in detecting functional diversity (Aglieri et al., 2021). In contrast, PAM was redundant in detecting functional diversity, detecting many small, arboreal taxa while omitting larger terrestrial species. This bias toward sound-producing taxa means functionally important mammals that vocalise infrequently remain undetected, potentially misrepresenting community functioning.

Trait-based approaches have their limitations. Many taxa lack published trait data, traits are often generalised across regions or life stages, and eDNA methods often fail to resolve taxa to species level (Condachou et al., 2023). While eDNA read counts or acoustic detection rates may correlate with abundance to some extent (Di Muri et al., 2020; Doser et al., 2021), these cannot currently estimate abundance reliably, limiting the use of metrics like RaoQ (Aglieri et al., 2021). These limitations, combined with no single method detecting the full range of functional traits, caution against overreliance on trait databases. However, acoustic and eDNA data can be archived and reanalysed as reference databases and classifiers improve (Condachou et al., 2023; Darras et al., 2025), offering future opportunities that could expand the functional scope of these methods. Our findings demonstrate that functional insights could be missed by using single methods in isolation.

### Trophic Complexity

Constructing complete trophic networks requires integrating methods that capture different components of community structure. PAM generated the most connected network, reflecting comprehensive coverage of bird taxa across trophic positions, but missed most mammalian interactions, which were primarily detected with eDNA. Camera traps contributed few interactions and were biased toward large, high-trophic-level species, consistent with known body size-trophic position relationships (Brose et al., 2019). Soil eDNA underperformed overall but uniquely detected *Accipiter gentilis*, the top predator. These method-specific biases reduced linkage density and robustness in single-method networks. Integrating methods proved critical: PAM and tree rolling together captured nearly 90% of observed links, rising to 98% when all eDNA substrates were included, demonstrating the value of method integration to avoid underestimation of trophic complexity.

Differences in temporal resolution of methods further influence inferred trophic interactions across methods. PAM and camera traps provide temporally explicit records (Sugai et al., 2019), enabling fine-scale inference of co-occurrence. By contrast, eDNA integrates signals over longer periods: days to weeks in water (Mauvisseau et al., 2022), weeks to months in soil (Guthrie et al., 2024), and likely intermediate durations on tree surfaces (Valentin et al., 2021). These broad temporal windows may dilute fine-scale patterns of interactions or inflate apparent co-occurrence by capturing species present at different times. This is especially relevant in rewilding landscapes, where species are often transient, and trophic interactions are dynamic (Torres et al., 2018). Understanding these temporal mismatches is essential for interpreting network structure.

## Conclusion

Integrating non-invasive monitoring methods provides better vertebrate biodiversity insights than single methods. PAM excels for taxonomic richness and trophic complexity, while eDNA captures broader functional diversity. However, eDNA substrate selection needs consideration. Soil and water substrates integrate signals from terrestrial, aquatic, and subterranean environments, while tree rolling detects arboreal species.

Method selection should align with restoration goals, prioritizing complementarity over redundancy. Combining PAM with one eDNA method provides sufficient coverage. For connectivity monitoring, combining PAM with soil sampling best detects spatial differences. For trophic complexity, combining PAM with tree rolling suits wooded habitats, while soil or water eDNA may better suit other contexts. Although not compared here, airborne eDNA methods also show promise for landscape-scale monitoring, capturing homogenised community signals (Clare et al., 2024) and should be considered in future research.

Around 1000 days of PAM alongside 50 tree rolling or water samples achieved optimal taxonomic coverage. However, single-timepoint eDNA sampling may have underestimated eDNA potential compared to continuous PAM data, and further method comparisons would benefit from temporal replication. Continuous PAM suits monitoring annual trends, while intensive eDNA sampling provides multitrophic snapshots of functional diversity. Multi-tool monitoring that combines taxonomic and functional diversity and trophic complexity will enable more comprehensive monitoring for ecosystem restoration and rewilding.

## Acknowledgements

We thank Emilia and Roger Leese for access to the Natural Capital Laboratory, Kristy Adaway for field assistance. This work was funded by the University of Hull REWILD research cluster.

## Data Availability Statement

Data, code, DNA sequences and media files are available via Zenodo https://doi.org/10.5281/zenodo.15723455, and GitHub repositories https://github.com/clarecowgill/Cowgill-et-al.-2025-Multi-tool-monitoring. DNA sampling and sequencing protocol is on protocols.io: dx.doi.org/10.17504/protocols.io.eq2lyqm8mvx9/v1.

## Author contributions

Clare Cowgill, Lori Lawson Handley, James D. J. Gilbert, and Ian Convery conceived the study. Clare Cowgill conducted eDNA sampling, lab work, and analyses. Acoustic and camera data formed part of Ian Convery and Volker B. Deecke ’s ongoing monitoring. Graham Sellers provided bioinformatics support. Clare Cowgill and Lori Lawson Handley led writing; Lori Lawson Handley and James D. J. Gilbert advised on analyses. All authors edited and approved the final manuscript. The authors declare no conflicts of interest.

## References

Aglieri, G., Baillie, C., Mariani, S., Cattano, C., Calò, A., Turco, G., Spatafora, D., Di Franco, A., … Milazzo, M. (2021). Environmental DNA effectively captures functional diversity of coastal fish communities. Molecular Ecology, 30(13), 3127–3139. doi:10.1111/mec.15661

Allen, M. C., Kwait, R., Vastano, A., Kisurin, A., Zoccolo, I., Jaffe, B. D., Angle, J. C., Maslo, B., & Lockwood, J. L. (2023). Sampling environmental DNA from trees and soil to detect cryptic arboreal mammals. Scientific Reports, 13(1), 180. doi:10.1038/s41598-023-27512-8

Botta-Dukát, Z. (2005). Rao’s quadratic entropy as a measure of functional diversity based on multiple traits. Journal of Vegetation Science: Official Organ of the International Association for Vegetation Science, 16(5), 533–540. doi:10.1111/j.1654-1103.2005.tb02393.x

Boyse, E., Robinson, K. P., Carr, I. M., Valsecchi, E., Beger, M., & Goodman, S. J. (2025). Inferring species interactions from co-occurrence networks with environmental DNA metabarcoding data in a coastal marine food web. Molecular Ecology, 34(7), e17701. doi:10.1111/mec.17701

Brose, U., Archambault, P., Barnes, A. D., Bersier, L.-F., Boy, T., Canning-Clode, J., Conti, E., Dias, M., … Iles, A. C. (2019). Predator traits determine food-web architecture across ecosystems. Nature Ecology & Evolution, 3(6), 919–927. doi:10.1038/s41559-019-0899-x

Cadotte, M. W., Carscadden, K., & Mirotchnick, N. (2011). Beyond species: functional diversity and the maintenance of ecological processes and services: Functional diversity in ecology and conservation. The Journal of Applied Ecology, 48(5), 1079–1087. doi:10.1111/j.1365-2664.2011.02048.x

Carraro, L., Mächler, E., Wüthrich, R., & Altermatt, F. (2020). Environmental DNA allows upscaling spatial patterns of biodiversity in freshwater ecosystems. Nature Communications, 11(1), 3585. doi:10.1038/s41467-020-17337-8

Carver, S., Convery, I., Hawkins, S., Beyers, R., Eagle, A., Kun, Z., Van Maanen, E., Cao, Y., … Soulé, M. (2021). Guiding principles for rewilding. Conservation Biology: The Journal of the Society for Conservation Biology, 35(6), 1882–1893. doi:10.1111/cobi.13730

Chen, S. (2023). Ultrafast one-pass FASTQ data preprocessing, quality control, and deduplication using fastp. iMeta, 2(2), e107. doi:10.1002/imt2.107

Clare, E. L., Economou, C. K., Bennett, F. J., Dyer, C. E., Adams, K., Mcrobie, B., & Littlefair. (2024). Airborne environmental DNA for terrestrial vertebrate community monitoring. Molecular Ecology Resources, 24(2), 253–266.

Condachou, C., Milhau, T., Murienne, J., Brosse, S., Villéger, S., Valentini, A., Dejean, T., & Mouillot, D. (2023). Inferring functional diversity from environmental DNA metabarcoding. *Environmental DNA (Hoboken*, N.J.), 5(5), 934–944. doi:10.1002/edn3.391

Cowgill, C., Gilbert, J. D. J., Convery, I., & Lawson Handley, L. (2025). Monitoring terrestrial rewilding with environmental DNA metabarcoding: a systematic review of current trends and recommendations. Frontiers in Conservation Science, 5. doi:10.3389/fcosc.2024.1473957

Darras, K. F. A., Rountree, R. A., Van Wilgenburg, S. L., Cord, A. F., Pitz, F., Chen, Y., Dong, L., Rocquencourt, A., … Wanger, T. C. (2025). Worldwide Soundscapes: A synthesis of Passive Acoustic Monitoring across realms. Global Ecology and Biogeography: A Journal of Macroecology, 34(5). doi:10.1111/geb.70021

de Bello, F., Carmona, C. P., Lepš, J., Szava-Kovats, R., & Pärtel, M. (2016). Functional diversity through the mean trait dissimilarity: resolving shortcomings with existing paradigms and algorithms. Oecologia, 180(4), 933–940. doi:10.1007/s00442-016-3546-0

Di Muri, C., Lawson Handley, L., Bean, C. W., Li, J., Peirson, G., Sellers, G. S., Walsh, K., Watson, H. V., … Hänfling, B. (2020). Read counts from environmental DNA (eDNA) metabarcoding reflect fish abundance and biomass in drained ponds. Metabarcoding and Metagenomics, 4. doi:10.3897/mbmg.4.56959

Doser, J. W., Finley, A. O., Weed, A. S., & Zipkin, E. F. (2021). Integrating automated acoustic vocalization data and point count surveys for estimation of bird abundance. Methods in Ecology and Evolution, 12(6), 1040–1049. doi:10.1111/2041-210x.13578

Dunne, J. A. (2005). The network structure of food webs. In Ecological Networks (pp. 27–92). Oxford University PressNew York, NY. doi:10.1093/oso/9780195188165.003.0002

Griffiths, N. P., Wright, R. M., Hänfling, B., Bolland, J. D., Drakou, K., Sellers, G. S., Zogaris, S., Tziortzis, I., … Vasquez, M. I. (2023). Integrating environmental DNA monitoring to inform eel (Anguilla anguilla) status in freshwaters at their easternmost range-A case study in Cyprus. Ecology and Evolution, 13(2), e9800. doi:10.1002/ece3.9800

Guthrie, A. M., Cooper, C. E., Bateman, P. W., van der Heyde, M., Allentoft, M. E., & Nevill, P. (2024). A quantitative analysis of vertebrate environmental DNA degradation in soil in response to time, UV light, and temperature. *Environmental DNA (Hoboken*, N.J.), 6(4). doi:10.1002/edn3.581

Harper, L. R., Lawson Handley, L., Hahn, C., Boonham, N., Rees, H. C., Gough, K. C., Lewis, E., Adams, I. P., … Hänfling, B. (2018). Needle in a haystack? A comparison of eDNA metabarcoding and targeted qPCR for detection of the great crested newt (Triturus cristatus). Ecology and Evolution, 8(12), 6330–6341. doi:10.1002/ece3.4013

Hart, E. E., Haigh, A., & Ciuti, S. (2023). A scoping review of the scientific evidence base for rewilding in Europe. Biological Conservation, 285(110243), 110243. doi:10.1016/j.biocon.2023.110243

Hartig, F., Abrego, N., Bush, A., Chase, J. M., Guillera-Arroita, G., Leibold, M. A., Ovaskainen, O., Pellissier, L., … Yu, D. W. (2024). Novel community data in ecology-properties and prospects. Trends in Ecology & Evolution, 39(3), 280–293. doi:10.1016/j.tree.2023.09.017

Hoefer, S., McKnight, D. T., Allen-Ankins, S., Nordberg, E. J., & Schwarzkopf, L. (2023). Passive acoustic monitoring in terrestrial vertebrates: a review. Bioacoustics, 32(5), 506–531. doi:10.1080/09524622.2023.2209052

Kahl, S., Wood, C. M., Eibl, M., & Klinck, H. (2021). BirdNET: A deep learning solution for avian diversity monitoring. Ecological Informatics, 61(101236), 101236. doi:10.1016/j.ecoinf.2021.101236

Kelly, R. P., Port, J. A., Yamahara, K. M., & Crowder, L. B. (2014). Using environmental DNA to census marine fishes in a large mesocosm. PloS One, 9(1), e86175. doi:10.1371/journal.pone.0086175

Leempoel, K., Hebert, T., & Hadly, E. A. (2020). A comparison of eDNA to camera trapping for assessment of terrestrial mammal diversity. *Proceedings*. Biological Sciences, 287(1918), 20192353. doi:10.1098/rspb.2019.2353

Mathews F., Kubasiewicz L. M.,Gurnell J., Harrower C. A., McDonald R. A., Shore R. F. (2018). Natural England Joint Publication JP025: A Review of the Population and Conservation Status of British Mammals.

Mauvisseau, Q., Harper, L. R., Sander, M., Hanner, R. H., Kleyer, H., & Deiner, K. (2022). The multiple states of environmental DNA and what is known about their persistence in aquatic environments. Environmental Science & Technology, 56(9), 5322–5333. doi:10.1021/acs.est.1c07638

McInerny, C. J., Musgrove, A. J., Gilroy, J. J., Dudley, S. P., Balmer, D., Batty, C., Crochet, P.-A., French, P., … the British Ornithologists’ Union Records Committee (BOURC). (2022). The British List: A Checklist of Birds of Britain (10th edition). The Ibis, 164(3), 860–910. doi:10.1111/ibi.13065

Mena, J. L., Yagui, H., Tejeda, V., Bonifaz, E., Bellemain, E., Valentini, A., Tobler, M. W., Sánchez-Vendizú, P., & Lyet, A. (2021). Environmental DNA metabarcoding as a useful tool for evaluating terrestrial mammal diversity in tropical forests. Ecological Applications: A Publication of the Ecological Society of America, 31(5), e02335. doi:10.1002/eap.2335

Newton, J. P., Allentoft, M. E., Bateman, P. W., van der Heyde, M., & Nevill, P. (2025). Targeting terrestrial vertebrates with eDNA: Trends, perspectives, and considerations for sampling. *Environmental DNA (Hoboken*, N.J.), 7(1). doi:10.1002/edn3.70056

Perino, A., Pereira, H. M., Navarro, L. M., Fernández, N., Bullock, J. M., Ceaușu, S., Cortés-Avizanda, A., van Klink, R., … Wheeler, H. C. (2019). Rewilding complex ecosystems. Science, 364(6438). doi:10.1126/science.aav5570

Pettorelli, N., & Bullock, J. M. (2023). Restore or rewild? Implementing complementary approaches to bend the curve on biodiversity loss. Ecological Solutions and Evidence, 4(2). doi:10.1002/2688-8319.12244

Poelen, J. H., Simons, J. D., & Mungall, C. J. (2014). Global biotic interactions: An open infrastructure to share and analyze species-interaction datasets. Ecological Informatics, 24, 148–159. doi:10.1016/j.ecoinf.2014.08.005

R Core Team. (2025) R: A Language and Environment for Statistical Computing. R Foundation for Statistical Computing. https://www.r-project.org/

Riaz, T., Shehzad, W., Viari, A., Pompanon, F., Taberlet, P., & Coissac, E. (2011). ecoPrimers: inference of new DNA barcode markers from whole genome sequence analysis. Nucleic Acids Research, 39(21), e145. doi:10.1093/nar/gkr732

Rognes, T., Flouri, T., Nichols, B., Quince, C., & Mahé, F. (2016). VSEARCH: a versatile open source tool for metagenomics. PeerJ, 4, e2584. doi:10.7717/peerj.2584

Ruppert, K. M., Kline, R. J., & Rahman, M. S. (2019). Past, present, and future perspectives of environmental DNA (eDNA) metabarcoding: A systematic review in methods, monitoring, and applications of global eDNA. GLOBAL ECOLOGY AND CONSERVATION, 17, e00547. doi:10.1016/j.gecco.2019.e00547

Ryan, E., Bateman, P., Fernandes, K., van der Heyde, M., & Nevill, P. (2022). eDNA metabarcoding of log hollow sediments and soils highlights the importance of substrate type, frequency of sampling and animal size, for vertebrate species detection. *Environmental DNA (Hoboken*, N.J.), 4(4), 940–953. doi:10.1002/edn3.306

Sales, N. G., McKenzie, M. B., Drake, J., Harper, L. R., Browett, S. S., Coscia, I., Wangensteen, O. S., Baillie, C., … McDevitt, A. D. (2020). Fishing for mammals: Landscape-level monitoring of terrestrial and semi-aquatic communities using eDNA from riverine systems. The Journal of Applied Ecology, 57(4), 707–716. doi:10.1111/1365-2664.13592

Sellers, G. S., Di Muri, C., Gómez, A., & Hänfling, B. (2018). Mu-DNA: a modular universal DNA extraction method adaptable for a wide range of sample types. Metabarcoding and Metagenomics, 2, e24556. doi:10.3897/mbmg.2.24556

Smith, M. D., Koerner, S. E., Knapp, A. K., Avolio, M. L., Chaves, F. A., Denton, E. M., Dietrich, J., Gibson, D. J., … Blair, J. M. (2020). Mass ratio effects underlie ecosystem responses to environmental change. The Journal of Ecology, 108(3), 855–864. doi:10.1111/1365-2745.13330

Sugai, L. S. M., Silva, T. S. F., Ribeiro, J. W., Jr, & Llusia, D. (2019). Terrestrial passive acoustic monitoring: Review and perspectives. Bioscience, 69(1), 15–25. doi:10.1093/biosci/biy147

Taberlet, P., Coissac, E., Pompanon, F., Brochmann, C., & Willerslev, E. (2012). Towards next-generation biodiversity assessment using DNA metabarcoding: NEXT-GENERATION DNA METABARCODING. Molecular Ecology, 21(8), 2045–2050. doi:10.1111/j.1365-294X.2012.05470.x

Torres, A., Fernández, N., Zu Ermgassen, S., Helmer, W., Revilla, E., Saavedra, D., Perino, A., Mimet, A., … Pereira, H. M. (2018). Measuring rewilding progress. Philosophical Transactions of the Royal Society of London. Series B, Biological Sciences, 373(1761). doi:10.1098/rstb.2017.0433

Valentin, R. E., Kyle, K. E., Allen, M. C., Welbourne, D. J., & Lockwood, J. L. (2021). The state, transport, and fate of aboveground terrestrial arthropod eDNA. *Environmental DNA (Hoboken*, N.J.), 3(6), 1081–1092. doi:10.1002/edn3.229

van der Heyde, M., Bunce, M., Wardell-Johnson, G., Fernandes, K., White, N. E., & Nevill, P. (2020). Testing multiple substrates for terrestrial biodiversity monitoring using environmental DNA metabarcoding. Molecular Ecology Resources, 20(3), 732–745. doi:10.1111/1755-0998.13148

Winiarska, D., Szymański, P., & Osiejuk, T. S. (2024). Detection ranges of forest bird vocalisations: guidelines for passive acoustic monitoring. Scientific Reports, 14(1), 894. doi:10.1038/s41598-024-51297-z

