## Supporting Information for "Multi-tool monitoring: integrating non-invasive methods to assess vertebrate diversity and trophic complexity in terrestrial rewilding"

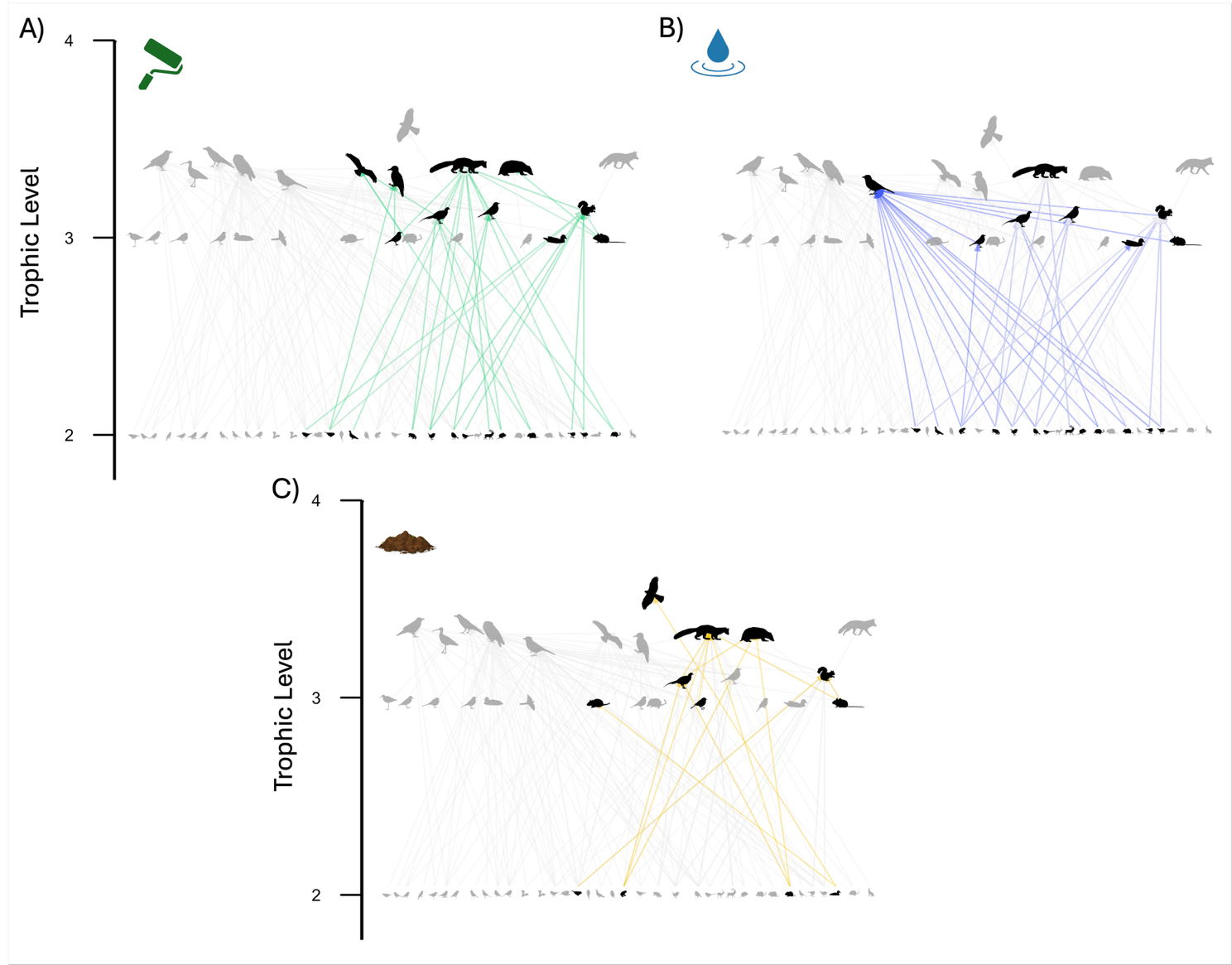
**Fig. S1:** Separate trophic co-occurrence networks for the eDNA sampling substrates: A) Tree rolling, B) Water, C) Soil.

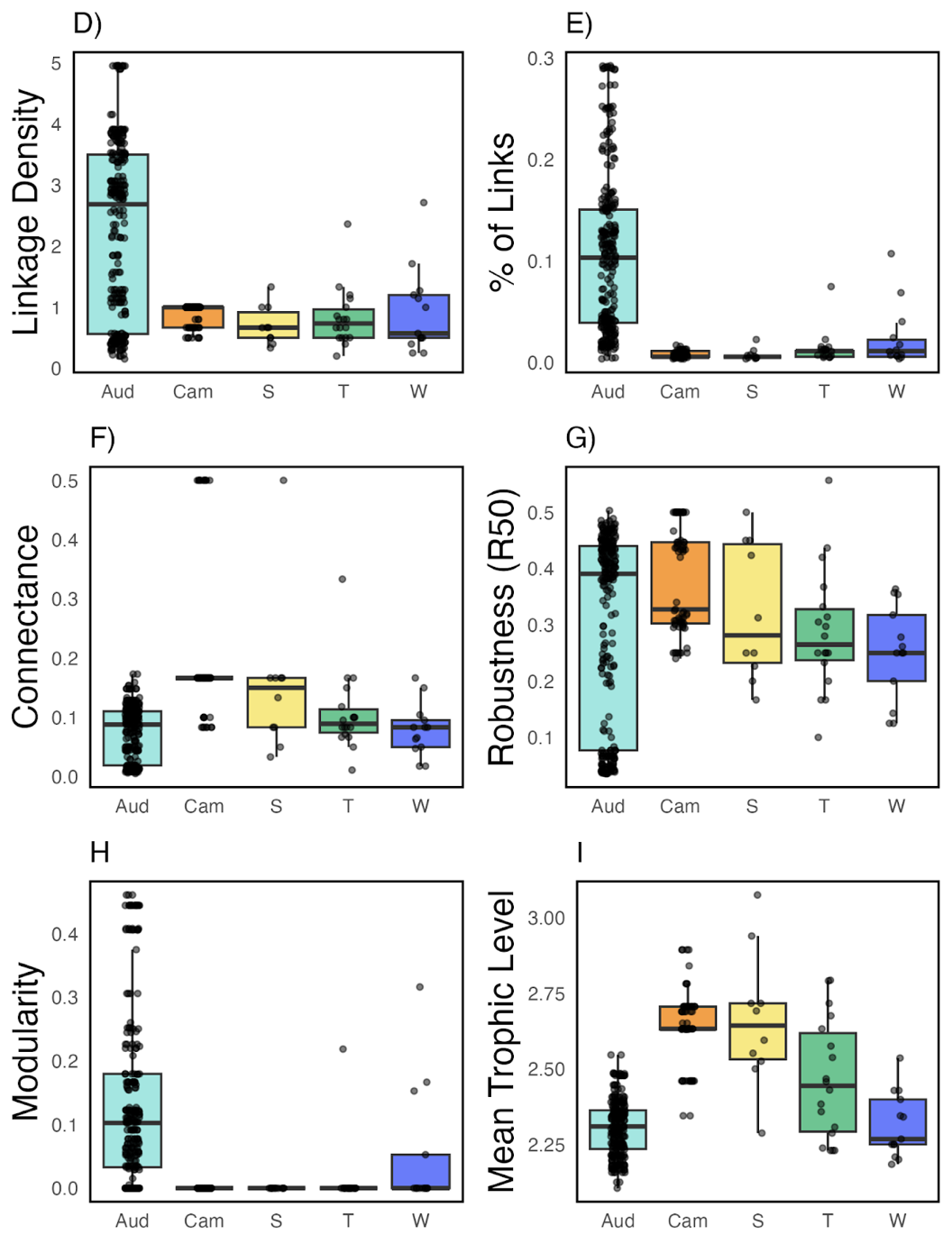
**Fig. S2:** Network metrics calculated for each method, shown by individual sample. *Note that camera and acoustic networks were generated using a rolling ±7-day detection window  (i.e. each point represents detections within a 14-day window centred on that time point), which results in non-independent observations across samples.*

**Supplementary Table 1:** Limits of Detection (LOD) read thresholds for each sample type and species combination where contamination was detected in corresponding blanks.

To account for potential contamination, we assessed blank control samples associated with each eDNA sample type (water, tree roller, soil, scat). We calculated species- and sample-type-specific limits of detection (LOD) using the mean and standard deviation of reads in corresponding field and laboratory blanks. For each species and sample type, we applied a conservative threshold by setting any read counts below the LOD to zero. Contamination was detected in blanks associated with multiple sample types and taxa.

| **Taxon** | **Sample Type** | **LOD Threshold (reads)** |
| --- | --- | --- |
| *Anatidae* | Tree roller | 81.3 |
| *Anatidae* | Water | 11.5 |
| *Anatidae* | Soil | 4.5 |
| *Anatidae* | Scat | 762.5 |
| *Arvicola amphibius* | Tree roller | 5.6 |
| *Arvicola amphibius* | Water | 8.4 |
| *Arvicola amphibius* | Soil | 1.7 |
| *Cervidae* | Tree roller | 727.7 |
| *Cervidae* | Soil | 449.7 |
| *Cervus elaphus* | Soil | 1017.5 |
| *Lissotriton helveticus* | Tree roller | 653.8 |
| *Lissotriton helveticus* | Water | 613.9 |
| *Lissotriton helveticus* | Soil | 199.3 |
| *Gallinula chloropus* | Tree roller | 7.0 |
| *Gallinula chloropus* | Water | 18.1 |
| *Gallinula chloropus* | Scat | 47.8 |
| *Martes martes* | Scat | 331.9 |
| *Rattus norvegicus* | Tree roller | 14.0 |
| *Rattus norvegicus* | Water | 5.6 |
| *Microtus agrestis* | Tree roller | 761.1 |
| *Microtus agrestis* | Water | 120.8 |
| *Phasianidae* | Tree roller | 727.7 |
| *Phasianidae* | Soil | 3319.5 |
| *Phasianidae* | Scat | 437.6 |
| *Rana temporaria* | Water | 2.5 |
| *Troglodytes troglodytes* | Water | 24.2 |

**Supplementary Table 2:** Taxonomic consolidation in the dataset for analyses across different methods.

| Taxon | Number of records | | | Taxon name after consolidation | Explanation |
| --- | --- | --- | --- | --- | --- |
|  | **Acoustic** | **Camera** | **eDNA** |  |  |
| Anas carolinensis |  |  | 1 | Anatidae | Since multiple Anatidae taxa were detected, for functional and trophic analyses, an average body mass of the detected taxa and the most common traits across species were used. |
| Aythya |  |  | 1 |  |  |
| Anatidae | 1 |  | 23 |  |  |
| Cervidae |  |  | 6 | Cervus nippon | All instances of Cervidae were in the same samples as Cervus nippon. Allowed us to keep distinct Cervus_elaphus detections |
| Cervus nippon |  | 99 | 73 |  |  |
| Columba livia |  |  | 5 | Columba | Since multiple Columba species were detected, for functional and trophic analyses, an average body mass of the detected taxa and the most common traits across species were used. |
| Columba oenas | 1 |  | 2 |  |  |
| Columba palumbus | 20 |  |  |  |  |
| Columba |  |  | 8 |  |  |
| Phasianus colchicus | 10 | 1 | 3 | Phasianidae | For functional and trophic analysis, Phasianus colchicus was used for all Phasianidae interactions. |
| Phasianidae |  |  | 16 |  |  |
| Fringilla coelebs | 31 |  |  | Fringilla | For functional and trophic analysis, Fringilla coelebs was used for all Fringilla interactions |
| Fringilla |  |  | 15 |  |  |
| Regulus regulus | 38 |  |  | Regulus | For functional and trophic analysis, Regulus regulus was used for all Regulus interactions |
| Regulus |  |  | 9 |  |  |
| Passer domesticus | 2 |  |  | Passer | For functional and trophic analysis, Passer domesticus was used for all Passer interactions |
| Passer |  |  | 1 |  |  |
| Phylloscopus trochilus | 31 |  |  | Phylloscopus | For functional and trophic analysis, Phylloscopus trochilus was used for all Phylloscopus interactions |
| Phylloscopus |  |  | 12 |  |  |

**Supplementary Table 3:** Presence and absence of each species for each method (shaded for presence), along with their functional traits used for analyses. Trophic niches: G (granivore), F (folivore), O (omnivore), I (invertivore), V (vertivore).

| ***Species*** | **Class** | **Acoustic** | **Camera** | **Soil** | **Scat** | **Tree rolling** | **Water** | **Lifestyle** | **Mass(g)** | **Trophic niche** |
| --- | --- | --- | --- | --- | --- | --- | --- | --- | --- | --- |
| *Bufo_bufo* | Amphibia |  |  |  |  |  |  | Semi-aquatic | 69.6 | I |
| *Lissotriton_helveticus* | Amphibia |  |  |  |  |  |  | Semi-aquatic | 1.8 | I |
| *Rana_temporaria* | Amphibia |  |  |  |  |  |  | Semi-aquatic | 22 | I |
| *Acanthis_cabaret* | Aves |  |  |  |  |  |  | Arboreal | 13.65 | I |
| *Accipiter_gentilis* | Aves |  |  |  |  |  |  | Semi-arboreal | 931.5 | V |
| *Actitis_hypoleucos* | Aves |  |  |  |  |  |  | Semi-aquatic | 48 | I |
| *Aegithalos_caudatus* | Aves |  |  |  |  |  |  | Arboreal | 8.5 | I |
| *Anatidae* | Aves |  |  |  |  |  |  | Semi-aquatic | 756.8 | O |
| *Anthus_pratensis* | Aves |  |  |  |  |  |  | Terrestrial | 18.5 | I |
| *Anthus_trivialis* | Aves |  |  |  |  |  |  | Semi-arboreal | 22.15 | I |
| *Buteo_buteo* | Aves |  |  |  |  |  |  | Semi-arboreal | 806.5 | V |
| *Carduelis_carduelis* | Aves |  |  |  |  |  |  | Arboreal | 15.45 | G |
| *Certhia_familiaris* | Aves |  |  |  |  |  |  | Arboreal | 9.1 | I |
| *Cinclus_cinclus* | Aves |  |  |  |  |  |  | Semi-aquatic | 60.25 | I |
| *Columba* | Aves |  |  |  |  |  |  | Terrestrial | 313 | G |
| *Corvus_corax* | Aves |  |  |  |  |  |  | Terrestrial | 1217.5 | O |
| *Corvus_corone* | Aves |  |  |  |  |  |  | Terrestrial | 526.85 | O |
| *Cuculus_canorus* | Aves |  |  |  |  |  |  | Arboreal | 111 | I |
| *Curruca_communis* | Aves |  |  |  |  |  |  | Semi-arboreal | 14.75 | I |
| *Cyanistes_caeruleus* | Aves |  |  |  |  |  |  | Arboreal | 11.5 | I |
| *Dendrocopos_major* | Aves |  |  |  |  |  |  | Arboreal | 74 | I |
| *Erithacus_rubecula* | Aves |  |  |  |  |  |  | Semi-arboreal | 17.55 | I |
| *Fringillidae* | Aves |  |  |  |  |  |  | Arboreal | 22.15 | I |
| *Fulica_atra* | Aves |  |  |  |  |  |  | Semi-aquatic | 777 | O |
| *Gallinula_chloropus* | Aves |  |  |  |  |  |  | Semi-aquatic | 348.5 | O |
| *Garrulus_glandarius* | Aves |  |  |  |  |  |  | Semi-arboreal | 164.4 | O |
| *Haematopus_ostralegus* | Aves |  |  |  |  |  |  | Semi-aquatic | 510 | I |
| *Hirundo_rustica* | Aves |  |  |  |  |  |  | Volant | 18.95 | I |
| *Lagopus_muta* | Aves |  |  |  |  |  |  | Terrestrial | 449 | G |
| *Lophophanes_cristatus* | Aves |  |  |  |  |  |  | Arboreal | 11.35 | I |
| *Loxia_curvirostra* | Aves |  |  |  |  |  |  | Arboreal | 41.7 | G |
| *Motacilla_alba* | Aves |  |  |  |  |  |  | Terrestrial | 20.5 | I |
| *Motacilla_cinerea* | Aves |  |  |  |  |  |  | Semi-aquatic | 17.85 | I |
| *Muscicapa_striata* | Aves |  |  |  |  |  |  | Arboreal | 16.3 | I |
| *Numenius_arquata* | Aves |  |  |  |  |  |  | Terrestrial | 725 | I |
| *Parus_major* | Aves |  |  |  |  |  |  | Arboreal | 18.25 | I |
| *Passer_domesticus* | Aves |  |  |  |  |  |  | Semi-arboreal | 29.25 | G |
| *Periparus_ater* | Aves |  |  |  |  |  |  | Arboreal | 9.8 | I |
| *Phasianidae* | Aves |  |  |  |  |  |  | Terrestrial | 1134 | O |
| *Phylloscopus_trochilus* | Aves |  |  |  |  |  |  | Arboreal | 9.1 | CI |
| *Plectrophenax_nivalis* | Aves |  |  |  |  |  |  | Terrestrial | 37.35 | G |
| *Pluvialis_apricaria* | Aves |  |  |  |  |  |  | Terrestrial | 206 | I |
| *Prunella_modularis* | Aves |  |  |  |  |  |  | Arboreal | 21.25 | I |
| *Pyrrhula_pyrrhula* | Aves |  |  |  |  |  |  | Arboreal | 31 | G |
| *Regulus_regulus* | Aves |  |  |  |  |  |  | Arboreal | 5.4 | I |
| *Saxicola_rubetra* | Aves |  |  |  |  |  |  | Semi-arboreal | 16.6 | I |
| *Scolopax_rusticola* | Aves |  |  |  |  |  |  | Terrestrial | 312 | I |
| *Sitta_europaea* | Aves |  |  |  |  |  |  | Arboreal | 23.05 | I |
| *Spinus_spinus* | Aves |  |  |  |  |  |  | Arboreal | 13.8 | G |
| *Strix_aluco* | Aves |  |  |  |  |  |  | Arboreal | 495.5 | V |
| *Sylvia_atricapilla* | Aves |  |  |  |  |  |  | Arboreal | 18.6 | I |
| *Tachybaptus_ruficollis* | Aves |  |  |  |  |  |  | Semi-aquatic | 135 | I |
| *Tringa_nebularia* | Aves |  |  |  |  |  |  | Semi-aquatic | 187 | I |
| *Troglodytes_troglodytes* | Aves |  |  |  |  |  |  | Arboreal | 8.9 | I |
| *Turdus_merula* | Aves |  |  |  |  |  |  | Semi-arboreal | 97 | I |
| *Turdus_philomelos* | Aves |  |  |  |  |  |  | Semi-arboreal | 76 | I |
| *Turdus_viscivorus* | Aves |  |  |  |  |  |  | Semi-arboreal | 117.7 | I |
| *Apodemus_sylvaticus* | Mammalia |  |  |  |  |  |  | Terrestrial | 21.9 | O |
| *Arvicola_amphibius* | Mammalia |  |  |  |  |  |  | Semi-aquatic | 120 | G |
| *Capreolus_capreolus* | Mammalia |  |  |  |  |  |  | Terrestrial | 22502.01 | F |
| *Cervus_elaphus* | Mammalia |  |  |  |  |  |  | Terrestrial | 79000 | F |
| *Cervus_nippon* | Mammalia |  |  |  |  |  |  | Terrestrial | 52999.99 | F |
| *Lepus_timidus* | Mammalia |  |  |  |  |  |  | Terrestrial | 3105.42 | F |
| *Martes_martes* | Mammalia |  |  |  |  |  |  | Semi-arboreal | 1299.99 | O |
| *Meles_meles* | Mammalia |  |  |  |  |  |  | Terrestrial | 11884.03 | O |
| *Microtus_agrestis* | Mammalia |  |  |  |  |  |  | Terrestrial | 35.87 | G |
| *Myodes_glareolus* | Mammalia |  |  |  |  |  |  | Terrestrial | 20.73 | O |
| *Myotis_daubentonii* | Mammalia |  |  |  |  |  |  | Volant | 7.63 | I |
| *Neomys_fodiens* | Mammalia |  |  |  |  |  |  | Semi-aquatic | 15.26 | I |
| *Oryctolagus_cuniculus* | Mammalia |  |  |  |  |  |  | Terrestrial | 1590.57 | F |
| *Pipistrellus_pipistrellus* | Mammalia |  |  |  |  |  |  | Volant | 5.3 | I |
| *Pipistrellus_pygmaeus* | Mammalia |  |  |  |  |  |  | Volant | 5 | I |
| *Rattus_norvegicus* | Mammalia |  |  |  |  |  |  | Terrestrial | 282.89 | O |
| *Sciurus_carolinensis* | Mammalia |  |  |  |  |  |  | Semi-arboreal | 545.4 | G |
| *Sciurus_vulgaris* | Mammalia |  |  |  |  |  |  | Semi-arboreal | 333 | G |
| *Sorex_araneus* | Mammalia |  |  |  |  |  |  | Terrestrial | 9.18 | I |
| *Sorex_minutus* | Mammalia |  |  |  |  |  |  | Terrestrial | 4.32 | I |
| *Talpa_europaea* | Mammalia |  |  |  |  |  |  | Fossorial | 87.53 | I |
| *Vulpes_vulpes* | Mammalia |  |  |  |  |  |  | Terrestrial | 4820.36 | O |

**Supplementary Table 4:** Kruskal Wallis and Dunn test results of species richness. P values adjusted with Benjamini-Hochberg.

| **Species richness: Kruskal Wallis (X2 = 197.9, p < 0.001)** | | | | | |
| --- | --- | --- | --- | --- | --- |
|  | PAM | Cameras | Scat | Soil | Tree |
| Camera | **12.38, 0.000** |  |  |  |  |
| Scat | **4.80, 0.000** | **-1.87, 0.035** |  |  |  |
| Soil | **9.56, 0.000** | **-3.17, 0.001** | 0.36, 0.358 |  |  |
| Tree rolling | **3.57, 0.000** | **-6.60, 0.000** | **-2.22, 0.016** | **-4.25, 0.000** |  |
| Water | **3.62, 0.000** | **-7.35, 0.000** | **-2.40, 0.011** | **-4.81, 0.000** | -0.16, 0.435 |

**Supplementary Table 5:** Pairwise PERMANOVA results for the community composition detected with each monitoring method using Jaccard dissimilarity. R² values first, with adjusted p value (Bonferroni) after.

|  | Acoustic | Camera | Scat | Soil | Tree |
| --- | --- | --- | --- | --- | --- |
| Camera | **0.49, 0.015** |  |  |  |  |
| Scat | **0.30, 0.015** | **0.24, 0.015** |  |  |  |
| Soil | **0.22, 0.015** | **0.19, 0.015** | **0.07, 0.015** |  |  |
| Tree | **0.20, 0.015** | **0.23, 0.015** | **0.16, 0.015** | **0.05, 0.015** |  |
| Water | **0.27, 0.015** | **0.20, 0.015** | **0.15, 0.015** | **0.06, 0.015** | **0.09, 0.015** |

**Supplementary Table 6:** PERMANOVA results for the community composition detected within each habitat for each monitoring method using Jaccard dissimilarity.

| **PERMANOVA** | | | | | |
| --- | --- | --- | --- | --- | --- |
| **Method** | **Df** | **SumOfSqs** | **R2** | **F** | **p.value** |
| Aud | 3 | 2.890 | 0.318 | 6.366 | < 0.001 |
| W | 3 | 1.897 | 0.169 | 2.097 | < 0.001 |
| S | 3 | 4.052 | 0.134 | 3.706 | < 0.001 |
| Cam | 3 | 1.420 | 0.122 | 3.899 | < 0.001 |
| T | 1 | 0.768 | 0.074 | 2.063 | 0.007 |

| **Pairwise PERMANOVA: Camera** | | | | | | |
| --- | --- | --- | --- | --- | --- | --- |
|  | **Df** | **SumsOfSqs** | **F.Model** | **R²** | **p.value** | **p.adjusted** |
| Felled vs Conifer | 1 | 1.052 | 8.038 | 0.124 | 0.001 | **0.006** |
| Felled vs Broadleaf | 1 | 0.509 | 5.484 | 0.111 | 0.001 | **0.006** |
| Bog vs Broadleaf | 1 | 0.310 | 3.341 | 0.150 | 0.015 | 0.09 |
| Bog vs Conifer | 1 | 0.527 | 3.276 | 0.093 | 0.027 | 0.162 |
| Felled vs Bog | 1 | 0.033 | 0.522 | 0.014 | 0.83 | 1 |
| Conifer vs Broadleaf | 1 | 0.059 | 0.333 | 0.008 | 0.911 | 1 |
| **Pairwise PERMANOVA: Acoustics** | | | | | | |
|  | **Df** | **SumofSqs** | **F.Model** | **R²** | **p.value** | **p.adjusted** |
| Bog vs Broadleaf | 1 | 0.77 | 6.61 | 0.36 | 0.001 | **0.006** |
| Felled vs Broadleaf | 1 | 1.02 | 6.26 | 0.29 | 0.001 | **0.006** |
| Felled vs Conifer | 1 | 1.71 | 10.31 | 0.26 | 0.001 | **0.006** |
| Bog vs Conifer | 1 | 1.09 | 7.56 | 0.23 | 0.001 | **0.006** |
| Felled vs Bog | 1 | 0.60 | 3.75 | 0.20 | 0.002 | **0.012** |
| Conifer vs Broadleaf | 1 | 0.29 | 1.99 | 0.07 | 0.047 | 0.282 |
| **Pairwise PERMANOVA: Soil** | | | | | | |
|  | **Df** | **SumofSqs** | **F.Model** | **R²** | **p.value** | **p.adjusted** |
| Broadleaf vs Felled | 1 | 1.92 | 5.59 | 0.13 | 0.001 | **0.006** |
| Broadleaf vs Bog | 1 | 1.29 | 3.43 | 0.12 | 0.003 | **0.018** |
| Conifer vs Felled | 1 | 2.14 | 5.97 | 0.11 | 0.001 | **0.006** |
| Conifer vs Bog | 1 | 1.15 | 2.99 | 0.08 | 0.003 | **0.018** |
| Broadleaf vs Conifer | 1 | 0.97 | 2.45 | 0.05 | 0.009 | 0.054 |
| Felled vs Bog | 1 | 0.25 | 0.80 | 0.03 | 0.541 | 1 |
| **Pairwise PERMANOVA: Tree rolling** | | | | | | |
|  | **Df** | **SumofSqs** | **F.Model** | **R²** | **p.value** | **p.adjusted** |
| Conifer vs Broadleaf | 1 | 0.77 | 2.06 | 0.07 | 0.007 | **0.03** |
| **Pairwise PERMANOVA: Water** | | | | | | |
|  | **Df** | **SumofSqs** | **F.Model** | **R²** | **p.value** | **p.adjusted** |
| Conifer vs Felled | 1 | 1.09 | 4.18 | 0.24 | 0.001 | **0.006** |
| Conifer vs Bog | 1 | 0.70 | 2.29 | 0.08 | 0.005 | **0.03** |
| Felled vs Bog | 1 | 0.31 | 1.34 | 0.11 | 0.224 | 1 |
| Broadleaf vs Conifer | 1 | 0.60 | 1.77 | 0.08 | 0.032 | 0.192 |
| Broadleaf vs Felled | 1 | 0.67 | 2.40 | 0.29 | 0.036 | 0.216 |
| Broadleaf vs Bog | 1 | 0.42 | 1.27 | 0.07 | 0.236 | 1 |

**Supplementary Table 7:** Kruskal Wallis and Dunn test results of functional diversity metrics. P values adjusted with Bonferroni.

| **Rao’s Quadratic Entropy: Kruskal Wallis (X2 = 54.51, p < 0.001)** | | | | |
| --- | --- | --- | --- | --- |
|  | Acoustics | Cameras | Soil | Tree |
| Camera | -0.03, 1.000 |  |  |  |
| Soil | **-2.33, 0.000** | -1.11, 1.000 |  |  |
| Tree rolling | **-5.69, 0.000** | **-2.86, 0.021** | **-3.16, 0.008** |  |
| Water | **-6.13, 0.000** | **-2.96, 0.016** | **-3.41, 0.003** | -0.09, 1.000 |
| **Average Functional Redundancy: Kruskal Wallis (X2 = 88.40, p < 0.001)** | | | | |
|  | Acoustics | Cameras | Soil | Tree |
| Camera | **3.40, 0.003** |  |  |  |
| Soil | **7.34, 0.000** | 0.30, 1.000 |  |  |
| Tree rolling | **7.39, 0.000** | 0.49, 1.000 | 0.35, 1.000 |  |
| Water | **6.69, 0.000** | -0.05, 1.000 | -0.66, 1.000 | -0.98 1.000 |

**Supplementary Table 8:** AIC scores for each of the tested models for predicting taxonomic richness per sample. The model in bold was chosen as the best fitting model.

| **Model predictor** | **AIC** |
| --- | --- |
| Null | 1784.145 |
| ***Method type*** | ***1033.617*** |
| Habitat type | 1748.634 |
| Method + Habitat | 1029.403 |
| Method * Habitat | 1024.540 |

*The Poisson regression model using only method as a predictor sufficiently explained taxonomic richness across the different eDNA substrates, cameras, and acoustic recorders, accounting for 78.52% of the detection variability. Including habitat, either as an additive or interaction term, did not improve model fit enough to warrant the added term. This confirms that differences in taxonomic richness can be attributed to sample type, supporting our use of method-specific diversity metrics throughout our analyses.*

**Supplementary Table 9:** Coefficients of the GLM predicting taxonomic richness based on sampling method type. Estimates represent the log difference in taxonomic richness compared to the reference method (Acoustics).

| Method | Estimate | St. Error | z value | P(>\|z\|) |
| --- | --- | --- | --- | --- |
| Acoustic (intercept) | 2.52 | 0.04 | 59.47 | < 0.001 |
| Camera | -2.14 | 0.10 | -21.02 | < 0.001 |
| Scat | -1.64 | 0.21 | -7.88 | < 0.001 |
| Soil | -1.69 | 0.09 | -19.46 | < 0.001 |
| Tree | -0.98 | 0.10 | -10.09 | < 0.001 |
| Water | -0.83 | 0.08 | -9.88 | < 0.001 |

**Supplementary Table 10:** The number of samples (either DNA samples or week of passive monitoring) which did not detect any terrestrial vertebrate taxa.

| **Method** | **Samples collected** | **Empty samples** |
| --- | --- | --- |
| Acoustics | 45 | 0 |
| Cameras | 105 | 24 (23%) |
| Scat | 10   (including lab replicates) | 0 |
| Soil | 100 (including lab replicates) | 24 (24%) |
| Tree rolling | 31   (including diluted samples) | 3 (10%) |
| Water | 35   (including diluted samples) | 0 |

**Supplementary Table 11**: R packages used and their citations

| **Package** | **Citation** |
| --- | --- |
| **tidyverse** | Wickham, H., Averick, M., Bryan, J., Chang, W., McGowan, L. D., François, R., … Yutani, H. (2019). Welcome to the tidyverse. *Journal of Open Source Software*, 4(43), 1686. https://doi.org/10.21105/joss.01686 |
| **ggplot2** | Wickham, H. (2016). *ggplot2: Elegant graphics for data analysis*. Springer. |
| **lubridate** | Grolemund, G., & Wickham, H. (2011). Dates and times made easy with lubridate. *Journal of Statistical Software*, 40(3), 1–25. https://doi.org/10.18637/jss.v040.i03 |
| **iNEXT** | Hsieh, T. C., Ma, K. H., & Chao, A. (2016). iNEXT: An R package for rarefaction and extrapolation of species diversity (Hill numbers). *Methods in Ecology and Evolution*, 7(12), 1451–1456. https://doi.org/10.1111/2041-210X.12613 |
| **vegan** | Oksanen, J., Blanchet, F. G., Friendly, M., Kindt, R., Legendre, P., McGlinn, D., … Wagner, H. (2020). *vegan: Community ecology package*. R package version 2.5-7. https://CRAN.R-project.org/package=vegan |
| **pairwiseAdonis** | Martinez Arbizu, P. (2020). pairwiseAdonis: Pairwise multilevel comparison using adonis. R package version 0.4.<https://github.com/pmartinezarbizu/pairwiseAdonis> |
| **SYNCSA** | Debastiani, V. J., & Pillar, V. D. (2012). SYNCSA - R tool for analysis of metacommunities based on functional traits and phylogeny of the community components. Bioinformatics, 28, 2067–2068. <https://doi.org/10.1093/bioinformatics/bts325> |
| **dunn.test** | Dinno, A. (2017). dunn.test: Dunn’s test of multiple comparisons using rank sums. R package version 1.3.5. |
| **cheddar** | Hudson, L. N., Emerson, R., Jenkins, G. B., Layer, K., Ledger, M. E., Pichler, D. E., … Woodward, G. (2013). Cheddar: An R package for ecological analyses of food webs. *Methods in Ecology and Evolution*, 4(5), 495–500. https://doi.org/10.1111/2041-210X.12010 |
| **igraph** | Csardi, G., & Nepusz, T. (2006). The igraph software package for complex network research. *InterJournal Complex Systems*, 1695.<https://igraph.org> |
| **MASS** | Venables, W.N. & Ripley, B.D. (2002) *Modern Applied Statistics with S. 4th edn. Springer, New York* |

**Supplementary Table 12**: Sources of functional trait data used to compile the trait database for analysis, including database names and references.

| **Package** | **Citation** |
| --- | --- |
| **EU Birds** | Storchová, L., & Hořák, D. (2018). Life‐history characteristics of European birds. Global Ecology and Biogeography: A Journal of Macroecology, 27(4), 400–406. doi:10.1111/geb.12709 |
| **AVONET** | Tobias, J. A., Sheard, C., … Schleuning, M. (2022). AVONET: morphological, ecological and geographical data for all birds. Ecology Letters, 25(3), 581–597. doi:10.1111/ele.13898 |
| **PanTHERIA** | Jones, K. E., Bielby, J., … Purvis, A. (2009). PanTHERIA: a species‐level database of life history, ecology, and geography of extant and recently extinct mammals: Ecological Archives E090-184. Ecology, 90(9), 2648–2648. doi:10.1890/08-1494.1 |
| **AmphiBIO** | Oliveira, B. F., São-Pedro, V. A., … Costa, G. C. (2017). AmphiBIO, a global database for amphibian ecological traits. Scientific Data, 4(1), 170123. doi:10.1038/sdata.2017.123 |
